# Nuclear Phase Separation Drives NPM1-mutant Acute Myeloid Leukemia

**DOI:** 10.1101/2025.05.23.655671

**Authors:** Gandhar K. Datar, Elmira Khabusheva, Archish Anand, Joshua Beale, Marwa Sadek, Chun-Wei Chen, Evdokiia Potolitsyna, Nayara Alcantara-Contessoto, Guangyuan Liu, Josephine De La Fuente, Christina Dollinger, Anna Guzman, Alejandra Martell, Katharina Wohlan, Abhishek Maiti, Nicholas J. Short, S. Stephen Yi, Vibeke Andresen, Bjørn Tore Gjertsen, Brunangelo Falini, Rachel E. Rau, Lorenzo Brunetti, Nidhi Sahni, Margaret A. Goodell, Joshua A. Riback

**Author notes:** Co-first authors (Gandhar K. Datar, Elmira Khabusheva).

## Abstract

During cancer development, mutations promote gene expression changes that cause transformation. Leukemia is frequently associated with aberrant *HOXA* expression driven by translocations in nucleoporin genes or *KMT2A*, and mutations in *NPM1*. How disparate mutations converge on this regulatory pathway is not understood. Here we demonstrate that mutant NPM1 (NPM1c) forms nuclear condensates in multiple human cell lines, mouse models, and primary patient samples. We show NPM1c phase separation is necessary and sufficient to coordinate the recruitment of NUP98 and KMT2A to condensates. Through extensive mutagenesis and pharmacological destabilization of phase separation, we find that NPM1c condensates are necessary for regulating gene expression, promoting *in vivo* expansion, and maintaining the undifferentiated leukemic state. Finally, we show that nucleoporin and KMT2A fusion proteins form condensates that are biophysically indistinguishable from NPM1c condensates. Together, these data define a new condensate underlying leukemias that we term coordinating bodies (C-bodies), and propose C-bodies as a therapeutic vulnerability.

## Introduction

Protein networks orchestrate cellular activities that encompass nearly all biological processes. Precise control of these networks requires spatial organization of proteins into discrete subcellular compartments. Such compartmentalization spans multiple scales, and includes the separation of cytoplasm and nucleus, membrane-bound organelles like mitochondria, and membrane-less structures – also termed biomolecular condensates – such as nucleoli and super-enhancer condensates^1^. Enrichment of proteins within subcellular compartments enables focused protein interactions critical for normal function. For example, the nucleolus directs active ribosomal biogenesis and is driven by essential factors such as Nucleophosmin (NPM1)^2–4^. Consequently, NPM1 mislocalization into the nucleoplasm is a well-established hallmark of cellular stress and is frequently indicative of cellular dysfunction^5,6^, demonstrating the close relationship between protein localization and function.

In the context of cancer, aberrant protein localization can disrupt normal cellular processes such as self-renewal or differentiation programs and activate oncogenic pathways that drive transformation. For example, normal tumor-suppressing mechanisms are inhibited when mutations cause abnormal cytoplasmic export of nuclear-acting proteins such as p53, Rb, and BRCA1^7^. Some oncogenic mutations, such as PML::RARA fusions, can alter protein localization and prevent the formation of biomolecular condensates such as PML bodies^8,9^. In contrast, other mutations can mislocalize proteins by precipitating the formation of new condensates, including those that stabilize signaling hubs or developmentally regulated transcriptional programs^10,11^. Notably, many groups have reported pathogenic condensates that annex a shared network of regulatory proteins in cancer, including chromatin-associated BRD4-NUT, ENL-driven transcriptional condensates, and NUP98 fusion proteins^12–16^. The emergence of new pathogenic condensates as a consequence of protein mislocalization is a compelling model in cancer^17,18^; however, demonstrating the connection between condensate stability and malignant transformation remains a critical challenge.

The formation of biomolecular condensates has emerged as a ubiquitous biophysical phenomenon that governs aspects of nearly all cellular activities, serving as bioreactors, sensors, signaling hubs, and more^1,10,19^. Nucleoli, PML nuclear bodies, and nuclear pore complexes are examples of biomolecular condensates that act as organizing centers to enrich the local concentration of specific proteins and facilitate distinct cellular activities. Broadly, the term condensate refers to liquid-like phases formed from biomolecules — similar to the physics underlying oil phase separating from water but with substantially higher complexity^20–22^. Specifically, much of the field has focused on the role of homotypic interactions – occurring between the same biomolecules – which notably involve intrinsically disordered regions (IDRs). In contrast, our studies and others have highlighted the importance of heterotypic interactions – occurring between different biomolecules – in driving condensates in cells^22,23^. Moreover, multivalency via oligomerization is a common feature of condensate proteins^24^. For example, the nucleolar protein NPM1 contains an N-terminal self-pentamerization domain, a large IDR with two alternating acidic and basic tracts, and an RNA binding domain (RBD). Each of these motifs is implicated in NPM1 phase separation *in vitro*, however only the pentamerization domain and RBD are critical for driving heterotypic phase transitions that form the nucleolus in cells^3,22,25,26^.

*NPM1* is the most frequently mutated gene in adult acute myeloid leukemias (AML), yet the mechanisms through which these mutations drive leukemic transformation remain unresolved^27,28^. Essentially all *NPM1* mutations result in a 4-base insertion in one allele of the C-terminal exon which causes a frameshift that generates a novel nuclear export sequence (NES) in mutant NPM1 (NPM1c)^29–31^. This NES is bound by XPO1, the primary exportin, resulting in NPM1c export to the cytoplasm^32^. Despite its eviction from the nucleolus and nucleus, NPM1c paradoxically drives a characteristic *HOXA/MEIS1* gene expression program essential for leukemia^33–37^. Knockout of mutant *NPM1* downregulates gene activation, leading to myeloid differentiation and cell growth arrest^35^. Unlike other AML subtypes that share this gene expression program – such as leukemias driven by nucleoporin (e.g., *NUP98*) fusions and *KMT2A* rearrangements – NPM1c does not have a clear association with chromatin regulation, or even nuclear localization^34^.

Over the last two decades, a number of contradictory models have been proposed to connect NPM1c localization and leukemic transformation, including cytoplasmic export of key nuclear proteins, loss of function of wild-type NPM1 (NPM1wt), and direct chromatin binding in the nucleus^36–39^. Whether NPM1c interacts with NPM1wt or is meaningfully found in the nucleus are ongoing questions that necessitate rigorous quantitative assessments. Robust data linking any proposed mechanism for NPM1c and its role in promoting disease in models of *NPM1*-mutant AML is lacking, slowing the development of new targeted therapies.

Furthermore, the connection between *NPM1*-mutant AML, and other subtypes of leukemias driven by nucleoporin and KMT2A oncofusions remains unclear. Notably, these leukemias often share therapeutic sensitivities to MENIN inhibitors^40–42^ and XPO1 inhibitors^35,43^, suggesting a common mechanism underlying leukemogenesis. A major challenge is to identify a common targetable factor, if any, across multiple disease subtypes.

In this study, we discovered that NPM1c forms nuclear condensates driven by heterotypic phase separation in models of *NPM1*-mutant AML and primary patient samples. We determined that NPM1c condensates are necessary and sufficient for the recruitment of proteins implicated in *HOXA/MEIS1* regulation, which together form a new biomolecular condensate we have termed coordinating bodies (or C-bodies). We show for the first time that C-bodies are necessary for leukemia cell survival, oncogene expression (*HOXA/MEIS1*), evading differentiation, and promoting *in vivo* expansion. Critically, we find that cytoplasmic localization of NPM1c is not sufficient for maintaining disease.

Looking towards other leukemia subtypes that share *HOXA/MEIS1* gene expression, we discovered that Nucleoporin and KMT2A fusion proteins phase separate in cells and recruit C-body associated proteins. Furthermore, we determine that condensates driven by NPM1c and oncofusions are all biophysically indistinguishable C-bodies. Together, our data provide unprecedented clarity into the dynamic role of NPM1c in AML, and suggest that C-bodies may be a unifying feature of multiple subtypes of leukemias. More broadly, this work highlights important biophysical principles that explain how distinct oncogenic condensates can be driven by relatively small genetic events. Finally, we provide an experimental framework to determine whether multiple reported pathogenic condensates may coalesce to establish a common targetable mesoscale structure.

## Results

### NPM1c forms nuclear condensates that are distinct from nucleoli

To examine the role of NPM1c in the cytoplasm or nucleus, we first sought to carefully quantify and compare the concentration of mutant and wild-type proteins in different cellular compartments. To assess localization of both proteins simultaneously within the same cell, we used CRISPR to introduce fluorescent proteins into the terminal exon of the wild-type and mutant alleles of *NPM1* in OCI-AML3 cells, a model of *NPM1*-mutant AML. NPM1c was largely present in the cytoplasm of each cell as previously described, while NPM1wt was primarily confined to the nucleolus, consistent with seminal works (**Figure 1A-B**)^27,35^. Strikingly, we also observed bright puncta in the nucleus in which NPM1c – but not NPM1wt – was highly enriched. NPM1c also decorated the nuclear lamina. Subcellular localization was independent of the specific tag (**Figures S1A-C**) and puncta were confirmed with immunostaining using antibodies targeting NPM1c (**Figure S1D**). These nuclear puncta, herein referred to as NPM1c condensates, are distinct from other common nuclear bodies (**Figure S1F**).

**Figure 1:**
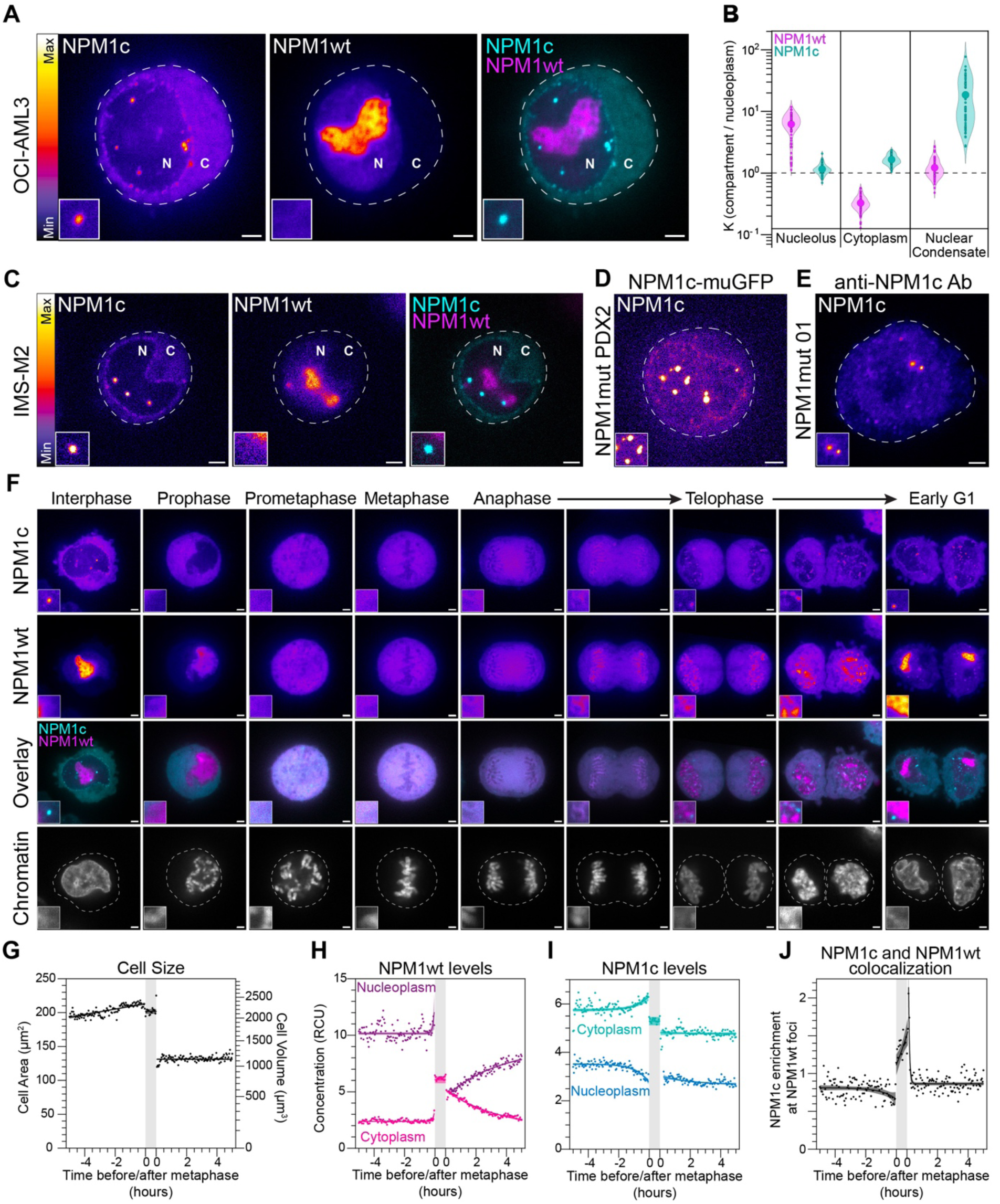
NPM1c resides in nuclear condensates that are established during cell division. **(A)** Live cell imaging of CRISPR-edited OCI-AML3 cells containing endogenous C-terminal-tagged NPM1c and NPM1wt (muGFP and mCherry respectively). **(B)** Quantification of partition coefficients (K) for NPM1wt (magenta) and NPM1c (cyan) within listed compartments relative to nucleoplasm (*n* = 100). **(C)** Live cell imaging of IMS-M2 cells with dual labeling as in panel A. **(D)** Live cell imaging of CRISPR-edited *NPM1*-mutant patient-derived xenograft (PDX2) cells with labeled NPM1c as in panel A. The window of minimum/maximum pixel intensities for this image was adjusted to optimally visualize condensates and the cytoplasm. **(E)** Immunostaining of primary NPM1-mutant AML patient sample (NPM1mut 01) with anti-NPM1c antibody. **(F)** Live cell imaging of dual-labeled OCI-AML3 cells stained with SPY650-DNA, a chromatin-labeling dye. Representative images of cell cycle stages are shown. **(G-K)** Quantification of live-cell imaging data of dual-labeled OCI-AML3 imaged every 3 minutes for ∼10hrs (*n* = 10). Metaphase (shaded gray) is defined by the average time between the earliest visualization of prometaphase and anaphase. **(G)** Quantification of cell area (µm^2^) and approximate volume (µm^3^). **(H)** NPM1wt and (**I**) NPM1c concentration in photons in relative concentration units (RCU). Timepoints reflecting prometaphase through anaphase have been annotated as cytoplasm. **(J)** Quantification of NPM1c enrichment in regions with high NPM1wt abundance. The brightest region (0.15µm^2^ box) of each cell containing NPM1wt was identified in individual cells and NPM1c level was measured. All images are shown in Fire LUT (see color scales in panels A & C) except for colocalization (cyan and magenta) and chromatin (grayscale). Dashed lines outline cytoplasmic membranes. N = nucleus. C = cytoplasm. White scale bars = 2µm. All insets are magnified 2µm square regions.

We next determined whether NPM1c condensates could be observed in other models of *NPM1*-mutant AML. Using the same knock-in strategy, we observed condensates in IMS-M2 cells – a separate *NPM1*-mutant cell line – and in a patient-derived xenograft (PDX2) model of *NPM1/FLT3*-mutant AML (**Figure 1C-D**)^35^. Furthermore, we performed immunostaining in patient samples using a primary *NPM1-*mutant leukemia and observed NPM1c condensates (**Figure 1E**). Notably these condensates were not observed in an NPM1^wt/wt^ primary AML sample (**Figure S1E**). These observations suggest that NPM1c condensates are a universal feature of *NPM1*-mutant leukemia.

To quantify the subcellular localization of NPM1c and NPM1wt, we utilized established microscopy methods that measure the strength of preferential recruitment or “transfer” between cellular microenvironments, here quantified as a ratio of levels known as the partition coefficient (K^tr^) (see Methods)^22,44^. As expected, NPM1wt strongly partitioned into nucleoli (K^tr^ = 6.2 ± 0.3) (**Figure 1B**). In contrast to long-held assumptions, NPM1c was only moderately enriched in the cytoplasm compared to the nucleoplasm (K^tr^ = 1.66 ± 0.03), but showed substantially higher partitioning (K^tr^ = 18 ± 2) into bright nuclear condensates (**Figure 1B**). Strikingly, these data reveal that NPM1c enrichment is heavily biased towards a previously unidentified nuclear condensate rather than the cytoplasm.

The stark separation of wild-type and mutant NPM1 into distinct condensates was surprising as both proteins retain an oligomerization domain that mediates NPM1 pentamerization. We therefore asked how this self-sorting was established in live cells.

### NPM1c condensates are established during cell division

During cell division, multiple cellular components, including the nuclear envelope and nucleolus, break down and reform. We hypothesized that this process would disrupt the separation of wild-type and mutant NPM1 and allow us to observe establishment of their compartmentalization. Therefore, we performed live imaging of these cells throughout the cell cycle.

NPM1wt recapitulated well-characterized behavior during mitosis^45,46^, allowing us to qualitatively prescribe stages of mitosis, including nucleolar breakdown during prophase, localization to the mitotic sheath in anaphase, and formation of pre-nucleolar bodies (PNBs) in telophase (**Figure 1F**).

Enrichment of NPM1c in the condensates and at the nuclear lamina was lost during prophase, consistent with the behavior of most nuclear bodies during mitosis^47^. After prophase and throughout anaphase, NPM1c showed strong colocalization with NPM1wt, including at the mitotic sheath. Strikingly, during telophase, NPM1c condensates reformed rapidly in tandem with NPM1wt pre-nucleolar bodies (PNBs), separating the mutant and wild-type NPM1 proteins (**Figure 1F**).

To better assess these dynamics we imaged at a higher frequency (every 3 minutes), identifying and segmenting individual cells undergoing mitosis. As expected, we observed an increase in cell size and volume as cells approached metaphase, a predictable halving of cell volume after division, and corresponding changes in the number of condensates per cell (**Figures 1G, S1G**).

As interphase cells approached metaphase, NPM1wt redistributed across cellular compartments during the last 6±2 minutes of prophase due to nucleolar breakdown (**Figure 1H**). Similarly, NPM1c enrichment in nuclear condensates and laminar puncta was lost during the 50±7 minutes of prophase and NPM1c concentration in the cytoplasm increased (**Figure 1I**). Surprisingly, NPM1c concentration decreased in the nucleoplasm, indicating that NPM1c-XPO1 complexes released from condensates and laminar puncta may be subsequently exported to the cytoplasm. After prophase, the localization of NPM1wt and NPM1c was largely indistinguishable (**Figure 1J**).

Following metaphase, NPM1wt and NPM1c rapidly segregate during early telophase (**Figure 1J**). After cytokinesis, NPM1wt levels are approximately equal in the nucleoplasm and cytoplasm, consistent with behavior in U2OS cells (**Figures S1H, S1J**). NPM1wt distribution across the nucleoplasm and cytoplasm gradually recapitulates the interphase state. Strikingly, NPM1wt in OCI-AML3 cells requires 3 hours (200±25 minutes) to establish its basal interphase level in the nucleoplasm, compared to only a half hour (25±1 minutes) in U2OS cells (**Figures 1H, S1J**). Indeed, expression of NPM1c in U2OS delays import of cytoplasmic NPM1wt (**Figure S1H-K**). In contrast, NPM1c levels in the cytoplasm did not significantly change following cytokinesis (**Figure 1I**). These results indicate that the presence of NPM1c substantially delays NPM1wt nuclear import.

Together, these data indicate two distinct sorting mechanisms for wild-type and mutant NPM1. While during metaphase they are found together, during telophase NPM1c and NPM1wt rapidly segregate into nuclear condensates and PNBs, respectively. Shortly thereafter, NPM1c appears trapped in nuclear condensates with little change in its nucleoplasmic and cytoplasmic levels. In contrast, NPM1wt continues slowly transiting from the cytoplasm into the nucleus. We anticipate this is due to the presence of mixed wild-type/mutant pentamers, which are rapidly exported via XPO1 binding. Over time, wild-type NPM1is able to complete its transit to the nucleolus without NPM1c.

### NPM1c condensates recruit factors implicated in gene regulation

Our findings suggest that NPM1c condensate formation is a highly regulated process, where exit from mitosis establishes condensates in the nucleus. Notably, the rapid reformation of NPM1c condensates distinct from PNBs indicates that interactions not native to NPM1wt likely drive condensate formation. During cell division, coordination of nucleoporins (NUPs) and exportins like XPO1 is necessary for nuclear envelope formation and chromosome segregation^48,49^. These processes largely coincide with NPM1c condensate formation. Therefore, we considered whether NUPs and XPO1 are enriched in NPM1c condensates.

We performed immunostaining in OCI-AML3 and IMS-M2 cells expressing endogenous GFP-labeled NPM1c (*NPM1*^WT/Degron^)^35^ for XPO1 and NUP98 - a core component of the nuclear pore complex. XPO1 and NUP98 formed puncta in the nucleus and at the nuclear periphery which qualitatively overlapped with NPM1c condensates (**Figures 2A and S2A**).To quantify colocalization of these puncta with NPM1c condensates, we calculated the radial distribution function (RDF) of relative immunostaining intensity around NPM1c condensates as previously described^50^. XPO1 and NUP98 show maximal spatial enrichment at the center of NPM1c condensates (i.e., zero displacement, **Figure 2B**). In contrast, COILIN staining for Cajal bodies – a well-known nuclear body – showed depletion from NPM1c condensates (**Figure 2A-B**).

**Figure 2:**
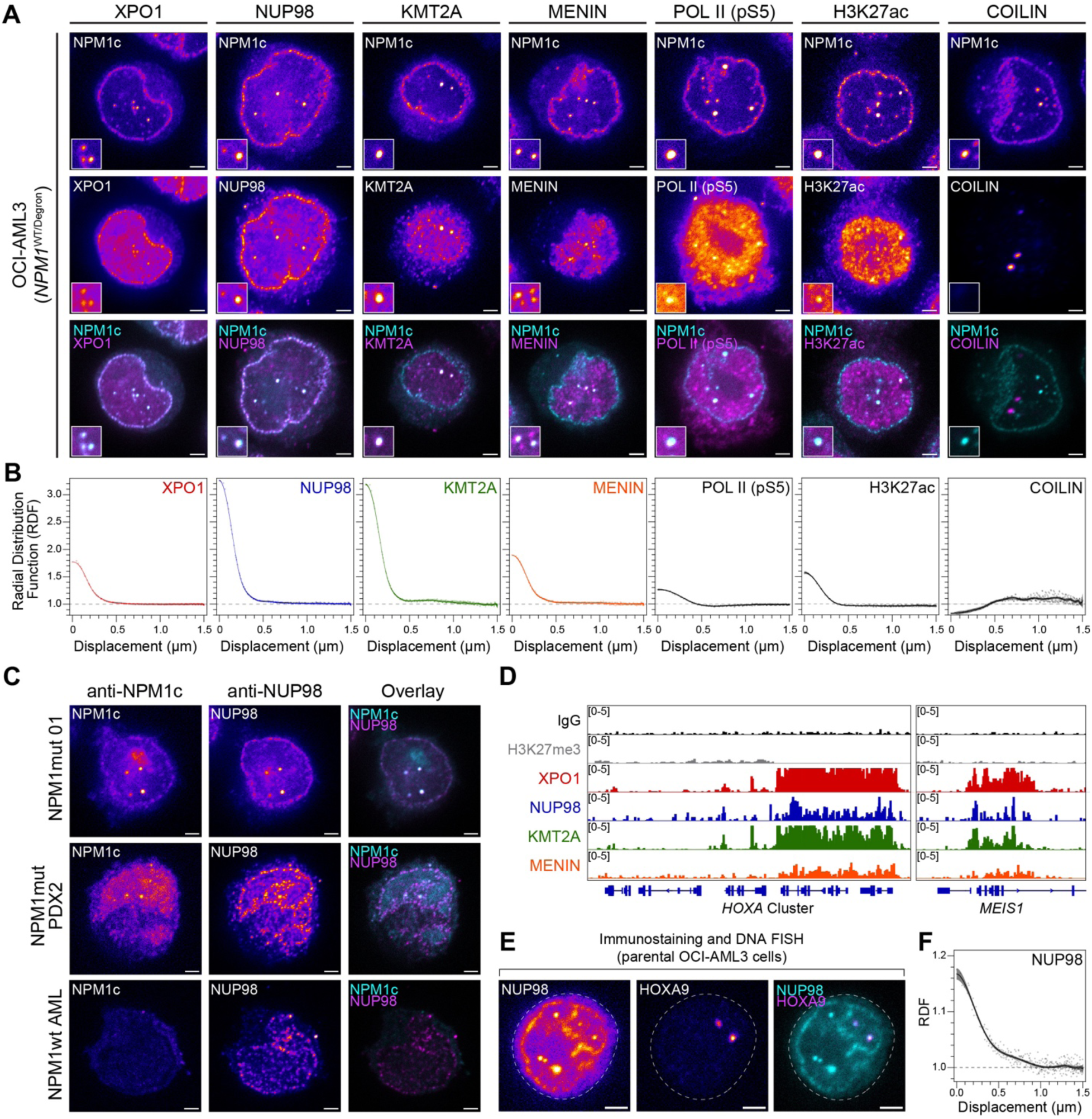
NPM1c condensates recruit chromatin regulatory proteins essential for leukemia. **(A)** Immunostaining of OCI-AML3 cells with GFP-tagged NPM1c (*NPM1*^WT/Degron^) for XPO1, NUP98, KMT2A, MENIN, and COILIN in fixed cells. **(B)** Radial distribution function (RDF) of antibody staining centered around the brightest nuclear NPM1c condensate per cell (*n* = 50 cells per staining). **(C)** Immunostaining of NPM1-mutant and wild-type NPM1 primary AML patient samples, and a PDX model of NPM1-mutant AML using anti-NPM1c and anti-NUP98 antibodies. **(D)** CUT&RUN sequencing data from *NPM1*^WT/Degron^ OCI-AML3 cells measuring chromatinenrichment of XPO1, NUP98, KMT2A, and MENIN at relevant genetic loci (1 replicate of *HOXA* gene cluster and *MEIS1* locus shown here, *n* = 2 replicates per antibody). IgG and H3K27me3 are displayed as negative and positive controls, respectively. All images are shown in Fire LUT except colocalization (cyan and magenta). **(E)** Combined immunostaining and DNA FISH of OCIAML3 cells using anti-NUP98 (cyan) antibodies and *HOXA9*-specific probes (magenta). **(F)** RDF of NUP98 antibody staining centered around *HOXA9* loci (*n* = 68 loci). White scale bars = 2μm. All insets are magnified 2μm square regions.

Recent reports have suggested that the histone methyltransferase KMT2A and its adaptor protein MENIN facilitate gene expression in tandem with NPM1c at characteristic *HOXA/MEIS1* genetic loci^36,37^. We therefore asked whether the KMT2A-MENIN complex enriches within NPM1c condensates. Similar to XPO1 and NUP98, KMT2A showed spatial enrichment in NPM1c condensates (**Figure 2A-B**). In contrast, MENIN formed dozens of puncta in the nucleoplasm, with many but not all overlapping with NPM1c condensates, consistent with its multifunctional role in transcription^51^ (**Figure 2A-B**). Inversely, all NPM1c condensates exhibited spatial enrichment of MENIN (**Figure 2B**). We also observed co-enrichment of NPM1c and NUP98 in primary AML patient samples (**Figure 2C**). These data suggest that NUP98, XPO1, KMT2A and MENIN interact together within NPM1c condensates, leading us to question if these condensate-associated proteins are enriched at *HOXA/MEIS1* loci.

To that end, we performed CUT&RUN sequencing in *NPM1*^WT/Degron^ OCI-AML3 cells to identify active chromatin regions bound by XPO1, NUP98, KMT2A, and MENIN. Consistent with previous reports^36,37^, we observed chromatin binding of XPO1, KMT2A, and MENIN at the *HOXA* cluster and *MEIS1*; we also identified NUP98 binding to these same loci (**Figure 2D**). Our data showed significant overlap with previously annotated NPM1c-bound genes identified by CHIP-seq in the same cell line,^36^ including genes in the *HOXA* cluster, *HOXB* cluster, and *MEIS1* **(Figure S3B, H).**

In light of these findings, we asked whether NPM1c condensates could be found at active chromatin regions. Immunostaining for active RNA polymerase II (POL II pS5) and histone marks associated with active enhancers (H3K27ac) revealed substantial co-localization with NPM1c condensates (**Figure 2A-B**). In order to examine whether condensates were localized to the *HOXA9* chromatin in intact nuclei, we performed DNA FISH. Notably, the anti-NPM1c antibody also marks nucleoli in OCI-AML3 cells, therefore we used NUP98 staining to unequivocally identify C-bodies across samples, given its high co-localization with NPM1c and presence at *HOX* loci by CUT&RUN. DNA FISH revealed that condensates are frequently associated with *HOXA9* chromatin (**Figure 2E-F**, **S3C**). Together these data suggest a strong correspondence between NPM1c condensates, the proteins enriched within them, and the genetic loci essential for driving *NPM1*-mutant AML; however, it remained unclear whether NPM1c was necessary for recruiting these proteins to nuclear condensates. Toward that end we next asked whether perturbation of NPM1c influenced condensate formation or the recruitment of associated proteins.

### NPM1c is necessary and sufficient for condensate formation and protein recruitment

To determine whether NPM1c is necessary for focal enrichment of XPO1, NUP98, KMT2A, and MENIN in the nucleus, we utilized an inducible degron model to degrade NPM1c (*NPM1*^WT/Degron^ cells)^35^. After treatment with dTAG-13, a small molecule that induces degradation, we observed an expected decrease in NPM1c abundance throughout the cell, along with increased expression of markers of myeloid differentiation, and loss of nuclear condensates (**Figures 3A-B, S3F-G**).

**Figure 3:**
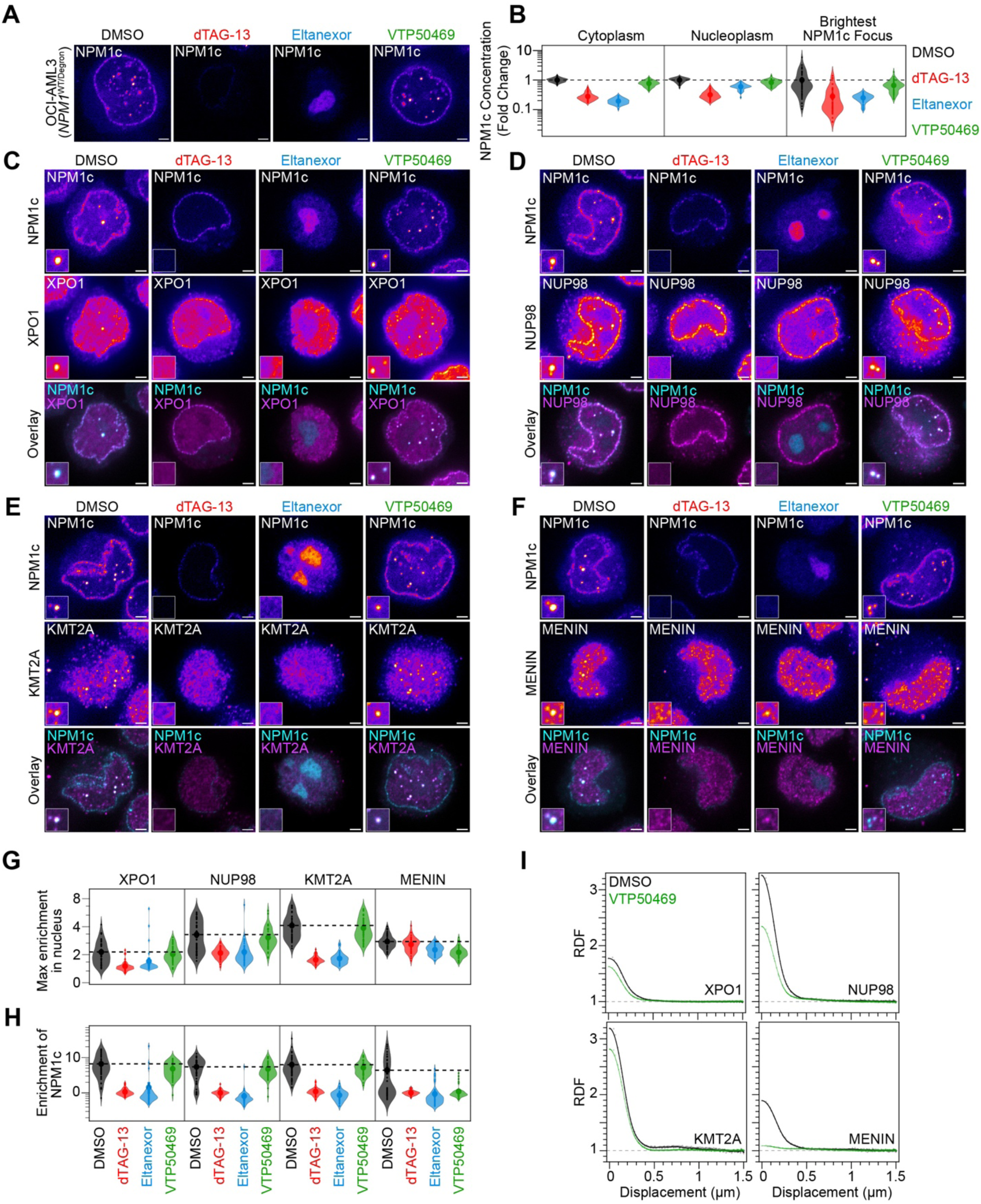
NPM1c condensates are necessary for protein recruitment. **(A)** Live cell imaging of *NPM1*^WT/Degron^ OCI-AML3 cells treated with DMSO, dTAG-13, Eltanexor, or VTP50469. **(B)** Quantification of NPM1c concentration across compartments after drug treatments (*n* = 50). **(C-F)** Immunostaining of *NPM1*^WT/Degron^ OCI-AML3 cells with antibodies targeting XPO1, NUP98, KMT2A, and MENIN after specific drug treatments. **(G)** Quantification of max protein enrichment in the nucleoplasm (*n* = 50). **(H)** Enrichment of NPM1c protein within max enriched ROI (*n* = 50). **(I)** RDF of XPO1, NUP98, KMT2A, and MENIN with DMSO or VTP50469 treatment, (*n* = 50). All cells were treated at same concentrations for 24hrs prior to live cell imaging or immunostaining (dTAG-13 = 500nM, Eltanexor = 100nM, VTP50469 = 300nM). All images are shown in Fire LUT except colocalization (cyan and magenta). White scale bars = 2µm. All insets are magnified 2µm square regions. For image analysis, the brightest region containing NPM1c or other protein in the nucleus is identified (0.15µm^2^ box) in individual cells and concentration (photons at reference settings) is measured and normalized to untreated controls.

To quantify changes in condensate loss, we identified the brightest region containing NPM1c in the nucleus (0.15µm^2^ box) in individual cells and measured NPM1c enrichment. We found that NPM1c-degradation resulted in loss of condensates compared to untreated cells (**Figures 3A-B**). We next asked whether condensate-associated proteins required NPM1c to form nuclear puncta. Immunostaining and quantification of the brightest focus per cell revealed a clear loss of XPO1, NUP98, and KMT2A nuclear puncta after NPM1c degradation. Notably, MENIN puncta persisted despite the loss of NPM1c (**Figures 3C-H**).

To test the sufficiency of NPM1c for condensate formation and protein recruitment in an NPM1wt context, we utilized U2OS cells. We expressed NPM1c in U2OS cells and observed *de novo* nuclear condensate formation (**Figure S2D-E**). Critically, this condensate formation did not rely on artificial genetic elements, such as LacO arrays, to induce NPM1c condensation^37^. In fact, we further validated *de novo* NPM1c condensate formation in the HL-60 acute promyelocytic leukemia cell line with overexpression and in human primary CD34+ cells derived from cord blood in which we converted *NPM1wt* to *NPM1c* using CRISPR editing (**Figure S2B-C**). As a result, we find that NPM1c is sufficient for condensate formation in hematologic and non-hematologic contexts.

We next asked whether NPM1c was sufficient to recruit proteins to nuclear condensates. Immunostaining of parental U2OS cells did not reveal XPO1, NUP98, or KMT2A nuclear puncta, however MENIN puncta were observed (**Figure S2D**). Strikingly, XPO1, NUP98, KMT2A, and MENIN were found within highly enriched nuclear puncta after NPM1c expression (**Figure S2E**). Consistent with OCI-AML3 cells (**Figure 2A)**, MENIN formed dozens of puncta in the nucleoplasm, with many but not all overlapping with NPM1c condensates. Together, these data indicate that NPM1c is both necessary and sufficient to coordinate the recruitment of XPO1, NUP98, KMT2A, and MENIN into previously unreported nuclear condensates, hereafter referred to as coordinating bodies (C-bodies).

### Multiple protein interactions are necessary for condensate stability, composition, and chromatin binding

We next asked whether specific protein-protein interactions were necessary for C-body formation. To probe these dependencies, we utilized well-established pharmacological strategies to block XPO1-NPM1c and MENIN-KMT2A interactions^35,42,52,53^. Importantly, XPO1 and MENIN inhibitors alter chromatin enrichment of specific proteins, halt essential leukemic transcriptional programs, and demonstrate clinical activity in *NPM1*-mutant AML patients^41,54^.

Consistent with previous work, treatment with the XPO1-inhibitors Eltanexor and Selinexor blocked NPM1c cytoplasmic export, and enhanced NPM1c localization to the nucleolus (**Figure 3A-B, S4A-D**)^35,53^. We also observed a concomitant decrease in *HOXA/MEIS1* expression and evidence of myeloid differentiation (**Figure S3E-G**). C-bodies were lost upon XPO1-inhibition, suggesting that the XPO1-NPM1c interaction is necessary for the condensates. Immunostaining revealed a loss of focal enrichments of XPO1, NUP98, and KMT2A in the nucleus (**Figure 3C-E, G-H, S4A-D**). In contrast, MENIN puncta persisted but were not enriched with NPM1c as seen in untreated cells (**Figure 3F-H, S4A-D**).

Next, we treated cells with the MENIN inhibitors VTP50469 and MI-503 and observed that MENIN-inhibition did not substantially impact NPM1c localization or C-body formation (**Figure 3A-B, S4A-D**). Immunostaining showed that XPO1, NUP98, and KMT2A persisted in nuclear puncta co-localizing with C-bodies (**Figure 3C-E, G-I, S4A-D**). In contrast, MENIN was specifically depleted from these condensates, despite forming numerous puncta elsewhere in the nucleus (**Figure 3F, G-I, S4A-E**). This focal loss of MENIN enrichment implies that KMT2A and MENIN interact within C-bodies, however this interaction is not necessary for condensate formation. Critically, MENIN depletion from condensates was associated with decreased *HOXA/MEIS1* expression and increased myeloid differentiation (**Figure S3E-G)**, suggesting that C-body composition is essential for its function in driving gene expression.

Our data demonstrate that degradation of NPM1c, inhibition of XPO1, or disruption of the MENIN-KMT2A interaction can alter C-body stability and composition. Notably, several groups have reported loss of chromatin enrichment of NPM1c, XPO1, KMT2A, and MENIN as a result of these treatment strategies^36,37,55,56^. Indeed, we observed decrease in XPO1, KMT2A, and MENIN binding at *HOXA/MEIS1* after dTAG-13, Eltanexor, or VTP50469 treatment that are consistent with previous reports (**Figure S3H**)^36,37^. We also report here that NUP98 recapitulates this pattern across drug treatments (**Figure S3H**). Together, our data suggest that C-bodies facilitate chromatin binding of gene regulatory proteins in *NPM1*-mutant AML.

### Pharmacological C-body disruption hinders *in vivo* models of NPM1c AML

Our data have established that C-bodies are highly enriched with KMT2A, MENIN, and other proteins which collectively bind active chromatin regions essential for leukemias (e.g. HOXA9). We further demonstrated that MENIN inhibition alters C-body composition and releases C-body-associated proteins from the chromatin. To establish the relationship between C-body composition, gene expression, and differentiation status, we turned to *in vivo* and *ex vivo* models of *NPM1*-mutant AML. Here, we applied MENIN inhibition to release C-body associated proteins from critical genes and examine the subsequent leukemia phenotype.

Using an inducible mouse model of *NPM1*-mutant pre-leukemia^57^, we observed C-body formation and focal co-enrichment of NUP98 and MENIN upon induction in murine HSPCs (**Figure 4A**). To determine the link between C-bodies, *HOXA/MEIS1* transcription, and cellular differentiation in a bonafide leukemia, we developed a rapid AML model utilizing *NRAS^G12D^*-GFP retroviral overexpression in *Dnmt3a*-mutant;*Npm1-*mutant murine HSPCs in line with other strategies^58^ (**Figure 4B and S6A**). We isolated GFP+ progenitors (c-Kit+;GFP+) from recipient mice with >80% donor chimerism in the peripheral blood and treated them with VTP50469 *ex vivo* for 24h to assess changes in C-body composition. While control-treated progenitors exhibited clear C-bodies, the MENIN inhibitor abolished association of MENIN from C-bodies marked by NUP98 (**Figure 4C**). RNA isolation after 72 hours revealed transcriptional downregulation of C-body-associated genes including *Hoxa9*, *Hoxa10*, *Meis1* (**Figure 4D**). After six days, cells treated with MENIN inhibitor showed significant signs of monocytic differentiation in contrast to untreated controls (**Figure 4E**). These data demonstrate that a mouse model of NPM1c-mutant leukemia exhibits clear C-bodies, the composition of which is critical for AML features.

**Figure 4:**
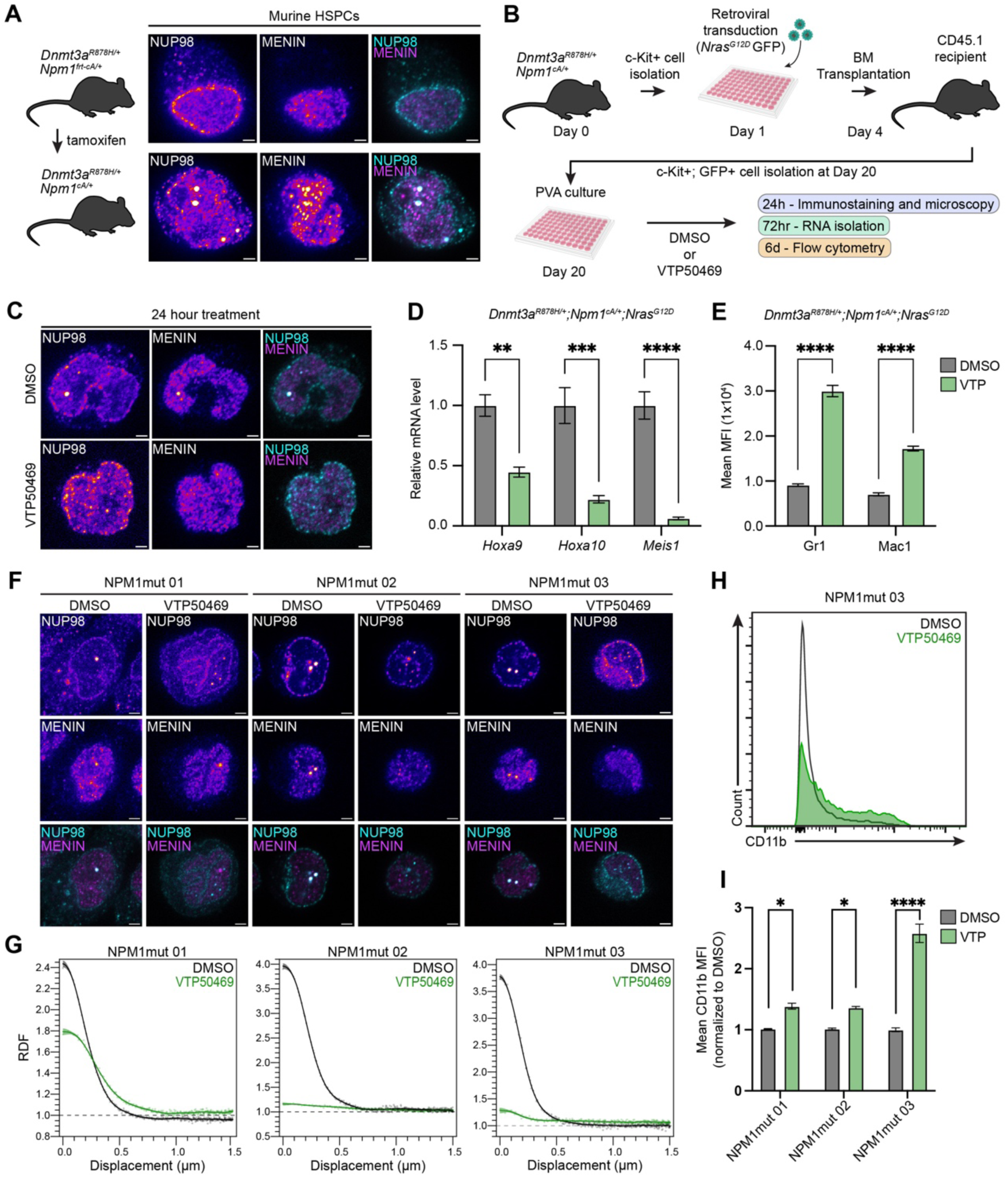
Compositional alteration of C-bodies impacts leukemic features in primary murine and human cells. **(A)** Npm1c induction with tamoxifen in mice (LEFT) and immunostaining of HSPCs isolated from NPM1frtC and NPM1c donor mice with anti-NUP98 (cyan) and anti-MENIN (magenta) antibodies. **(B)** Schematic representation of Npm1c AML model development for *ex vivo* studies. **(C)** Immunostaining of murine HSPCs treated with DMSO or VTP50469 *ex vivo* with anti-NUP98 (cyan) and anti-MENIN (magenta) antibodies. **(D)** Real-time qPCR analysis of *Hoxa9*, *Hoxa10* and *Meis1* mRNA levels and **(E)** flow cytometry analysis of myeloid differentiation markers Gr1 and Mac1 (MFI) in murine HSPCs treated with DMSO or VTP50469 *ex vivo*. **(F)** Immunostaining of human AML patient cells treated with DMSO or VTP50469 *ex vivo* with anti-NUP98 (cyan) and anti-MENIN (magenta) antibodies. **(G)** RDF of NUP98 and MENIN with DMSO or VTP50469 treatment (*n* = 20-60 cells per sample). **(H)** Representative histogram of CD11b fluorescence intensity in DMSO and VTP50469. **(I)** Fold change of mean CD11b fluorescence level in patient samples relative to DMSO cells. All cells were treated at the same concentrations for 24hrs prior to live cell imaging or immunostaining. Flow cytometry analysis was performed at day 3 and day 6 in human and murine cells respectively.

Next, we examined peripheral blood from *NPM1*-mutant AML patients with high blast counts (**Table S2**). Immunostaining of mononuclear cells revealed distinct C-bodies marked by focal co-enrichment of NUP98 and MENIN across all samples (**Figure 4F**). Treatment with VTP50469 led to a depletion of MENIN in C-bodies and a corresponding shift towards monocytic differentiation (**Figures 4F-I**), consistent with our observations in cell lines and murine HSPCs (**Figure 3, 4C**). Thus, we conclude that C-body composition is strongly associated with leukemogenic gene expression and differentiation in *NPM1*-mutant AML.

### Heterotypic phase separation drives C-body formation

Despite their disparate mechanisms of action, our data show that XPO1 and MENIN inhibitors alter the formation and composition of C-bodies, respectively. This convergent behavior on condensates containing several key leukemia-associated proteins suggests that these previously unreported structures may play an integral role in leukemogenesis. Thus, we turned our efforts to understanding the biophysical determinants of NPM1c condensate formation.

Phase separation is a concentration-dependent behavior defined by the presence of a phase boundary at a saturation concentration or *C*_sat_ ^20^. At concentrations above the *C*_sat_, condensate formation is favored. In multi-component phase separation, a stoichiometric imbalance in one component can alter condensate composition, size, and function. When heterotypic interactions drive phase separation, stoichiometric imbalance results in heterotypic destabilization^22^.

To this end, we asked whether C-bodies are phase-separated condensates, and if heterotypic interactions drive their formation. We examined their concentration-dependent behavior using a lentiviral system to overexpress full-length NPM1c (FL-NPM1c) in *NPM1*^WT/Degron^ OCI-AML3 cells. While low FL-NPM1c expression had no effect on endogenous C-bodies in the nucleus or enrichment at the nuclear periphery, high FL-NPM1c expression produced larger C-bodies, and reduced enrichment at the nuclear periphery, suggestive of heterotypic destabilization (**Figure S5A**). In contrast, overexpression of the fluorescent protein alone had no effect on endogenous C-bodies (**Figure S5C**). We next degraded endogenous NPM1c with dTAG-13 treatment, finding that only high-expressing FL-NPM1c cells retained C-body formation (**Figure S5B**). Overexpression of the fluorescent protein alone did not lead to C-body formation (**Figure S5D**). These data suggest a threshold expression level or *C_sat_* for FL-NPM1c in dTAG-13 treated cells (**Figure S5B**).

Heterotypic destabilization results in a non-fixed *C_sat_* that causes accumulation of excess molecules in the dilute phase^22^. For example, when nucleoli dissolve during prophase the nucleoplasm is flooded with excess NPM1wt (**Figure 1H**). Likewise, we hypothesized that destabilization of C-bodies would result in higher cytoplasmic accumulation of NPM1c, due to XPO1-mediated export of excess NPM1c released into the nucleoplasm. Consistent with our observations of NPM1c condensate loss in prophase (**Figure S1G**), overexpression of FL-NPM1c results in greater cytoplasmic export of endogenous NPM1c (**Figure S5E**). Altogether, these data indicate that C-bodies are driven by heterotypic phase separation in *NPM1*-mutant AML. Importantly, these results oppose a classical binding model that lacks phase separation, in which heterotypic destabilization due to protein over-expression should not be observed.

Finally, we examined whether heterotypic destabilization of C-bodies had an impact on cell growth and differentiation. Strikingly, cells overexpressing FL-NPM1c had reduced growth over time compared to cells expressing the fluorescent protein alone (**Figure S5F-G**), suggesting a decrease in cell fitness. Importantly, FL-NPM1c overexpression was sufficient to prevent cells from differentiating when treated with dTAG-13, implying that FL-NPM1c is functionally indistinguishable from endogenous NPM1c (**Figure S5G,K**). These data indicate that stable C-bodies, and the interactions within them, are important for leukemia maintenance. Furthermore, heterotypic destabilization of C-bodies decreased cell growth despite enhanced localization of endogenous NPM1c to the cytoplasm (**Figure S5F**). Altogether, these results establish C-bodies as phase separated compartments driven by heterotypic interactions whose stability dictates cell growth and the leukemic state.

### Heterotypic destabilization of C-bodies decreases cell growth

To better understand NPM1c’s role in condensate formation and cancer cell fitness, we focused on NPM1c protein architecture. The NPM1 protein has several regions including an oligomerization (self-pentamerization) domain, a conserved IDR with alternating acidic and basic stretches, and a C-terminal domain (CTD) that folds into a globular RNA-binding domain^26,59^. In the context of the common C-terminal AML mutation, the CTD is essentially unstructured^30^. To discern which domains contribute to C-body formation, we used a lentiviral system to express a series of truncated NPM1c proteins in *NPM1*^WT/Degron^ OCI-AML3 cells (**Figure 5A,B**).

**Figure 5:**
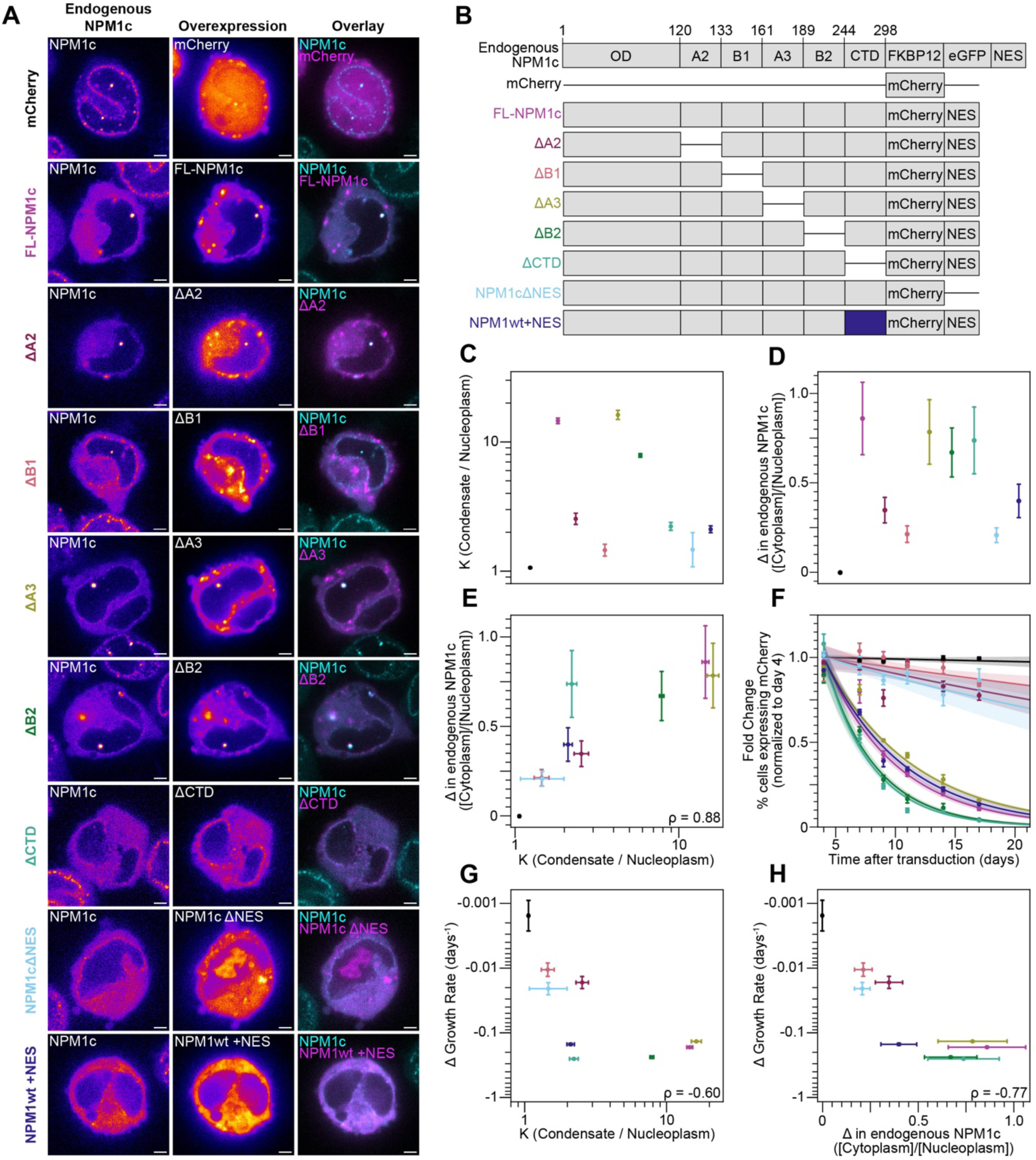
Destabilization of NPM1c condensates is associated with enhanced cytoplasmic export of NPM1c and reduced cell growth. **(A)** Live cell imaging of *NPM1*^WT/Degron^ OCI-AML3 cells (no dTAG treatment) expressing NPM1c truncations. **(B)** Schematic of NPM1c truncation series. **(C)** Partitioning (K) of each truncation into nuclear condensates (Condensate/Nucleoplasm). Error bars indicate fit parameter standard error. **(D)** Instantaneous change (as in Figure 4C) in endogenous NPM1c localization [Cytoplasm/Nucleoplasm] with expression of truncations. Error bars indicate fit parameter standard error. **(E)** Correlation between partitioning (K) and change in endogenous NPM1c localization. **(F)** Fold Change of % cells expressing truncations over 17 days (normalized to Day 4), measured by flow cytometry (includes repeated data for FL-NPM1c and mCherry as shown in Figure 4) *n* = 3 biological replicates per truncation. **(G)** Correlation of change in growth rate (shown as days^-1^) with partitioning (K) into C-bodies and **(H)** change in endogenous NPM1c localization [Cytoplasm/Nucleoplasm]. *n* > 40 for panels C-E, G, H. ρ is Spearman correlation coefficient. All images are shown in Fire LUT except colocalization (cyan and magenta). White scale bar = 2µm.

All proteins displayed clear cytoplasmic localization, and most enriched in C-bodies, including FL-NPM1c, ΔA2, ΔB1, ΔA3, ΔB2, ΔCTD, NPM1cΔNES, and NPM1wt+NES (**Figure 5A**). Truncated proteins exhibited a range of partition coefficients into C-bodies, and the fluorescent protein control (mCherry) was not enriched in condensates as expected (**Figure 5C**). Consistent with previous findings, NPM1cΔNES was also observed in the nucleolus (**Figure 5A, C**)^35^. We next asked how these truncated proteins altered endogenous NPM1c localization as in **Figure 4C**. All truncation variants enhanced cytoplasmic export of endogenous NPM1c; however, FL-NPM1c, ΔA3, ΔB2, and ΔCTD had a substantially greater effect on localization (**Figure 5D**). Indeed, the partitioning (K) of truncated proteins into C-bodies strongly correlated with cytoplasmic export of endogenous NPM1c (ρ = 0.88, **Figure 5E**). In this way, proteins that can favorably drive heterotypic phase separation destabilize C-bodies, resulting in displacement of endogenous NPM1c from condensates and enhanced export into the cytoplasm. These data suggest that multiple domains of NPM1c can contribute to heterotypic destabilization of endogenous condensates, in accordance with the multivalent interactions needed for phase separation.

As a result of these findings, we questioned if the degree of cytoplasmic export of endogenous NPM1c influenced cell growth as in **Figure 4D**. After expressing each truncated protein, we observed cell growth for approximately two weeks and identified two discrete behaviors (**Figure 5F**). Cells expressing ΔA2, ΔB1, and NPM1cΔNES had a minor decrease in growth compared to the mCherry control. In stark contrast, cells expressing FL-NPM1c, ΔA3, ΔB2, ΔCTD, and NPM1wt+NES had substantially decreased growth over time. Remarkably, we found that the quantitative change in growth rate was negatively correlated with partitioning into condensates and cytoplasmic export of endogenous NPM1c (ρ = -0.60 and -0.77, **Figure 5G, H**). Together, these data suggest that heterotypic destabilization of C-bodies impairs cell fitness.

### C-bodies are necessary for leukemia cell survival *in vitro* and *in vivo*

The results of the truncation study led us to ask whether any proteins capable of heterotypic destabilization could form condensates without endogenous NPM1c. Notably, the step-like response between partitioning and growth rate (Figure 5G) could indicate that ΔCTD and NPM1wt+NES, which have low partitioning but strong consequences for growth rate, remove interactions needed for phase separation and thus act as caps to destabilize C-bodies through loss of valence as seen in other contexts^24^. To this end, we expressed each truncation in *NPM1*^WT/Degron^ OCI-AML3 cells and treated them with dTAG-13 to degrade endogenous NPM1c. As shown previously here, FL-NPM1c recapitulated C-body formation, and fluorescent protein expression alone was not sufficient to form nuclear structures (**Figure S5A-B, 6A**). In addition, ΔA3, and ΔB2 formed nuclear condensates, suggesting that A3 and B2 domains are dispensable for phase separation. As expected, ΔCTD did not form nuclear condensates, indicating that the mutant-specific CTD is necessary for phase separation. ΔA2 and ΔB1 also lacked condensates, suggesting that a second interaction spanning the IDR of NPM1c is required. NPM1wt+NES did not form condensates, indicating that XPO1 binding to a C-terminal NES on NPM1wt is not sufficient for aberrant phase separation (Figure 6A). Finally, we did not observe C-body formation by NPM1cΔNES (Figure 6A). This protein maintains the full amino acid sequence of NPM1c and ostensibly all possible binding interactions, suggesting that it is a “C-body null” variant of NPM1c.

**Figure 6:**
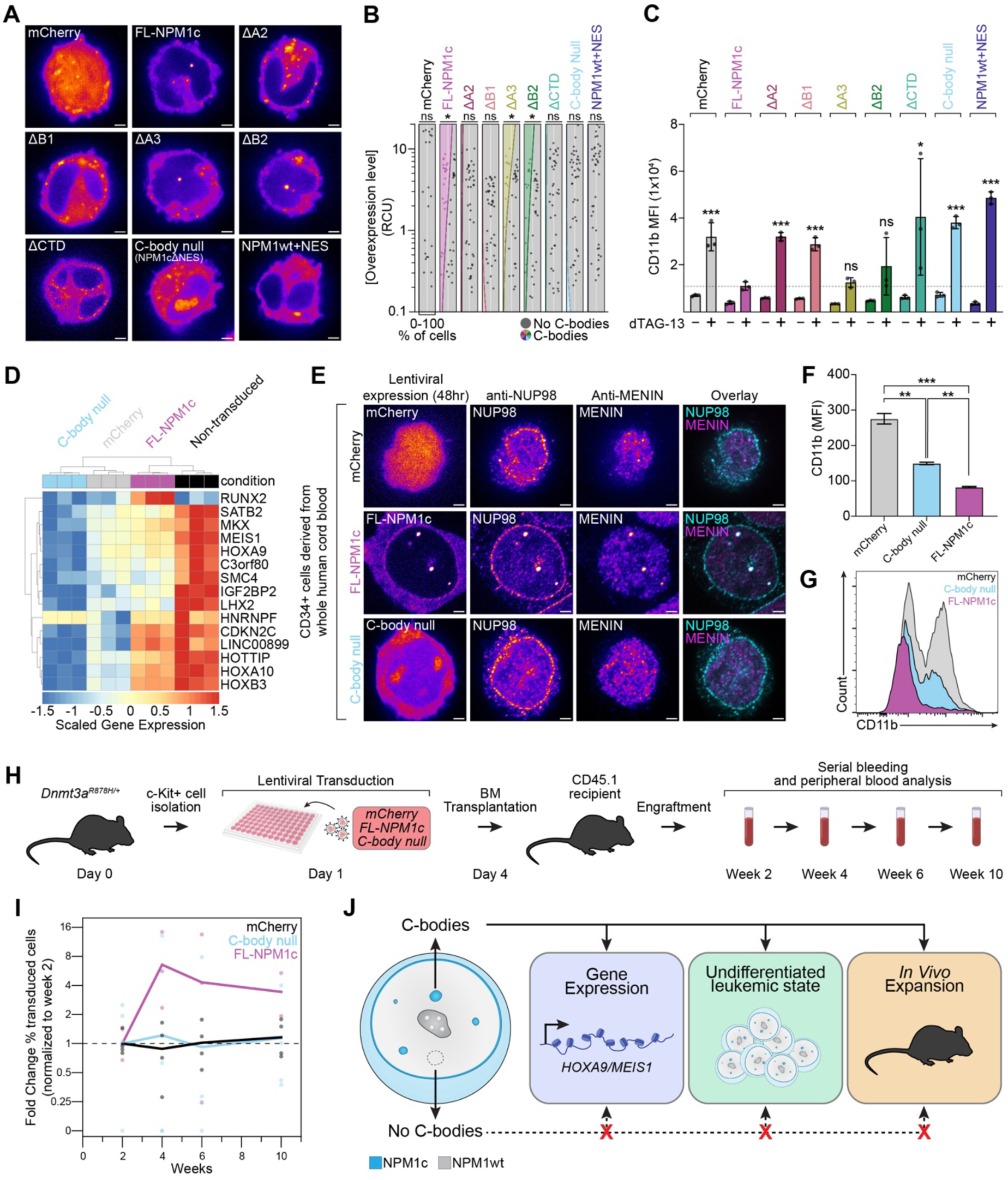
Multiple domains of NPM1c are required for condensate formation and maintaining the disease state. **(A)** Live cell imaging of *NPM1*^WT/Degron^ OCI-AML3 cells expressing NPM1c truncations after 7 days of dTAG-13 (500nM) treatment. **(B)** Quantification of NPM1c concentration in photons in relative concentration units (RCU) and condensate formation after dTAG-13 treatment (includes data repeated from Figure 4). Shaded regions represent fraction of cells without condensates (gray), or with condensates (multicolor). Spearman Rank Test, *p<.05, *n* > 40. **(C)** Median Fluorescence Intensity (MFI) of CD11b via flow cytometry after 14 days of DMSO or dTAG-13 treatment, *n* = 3 biological replicates per treatment and truncation. (includes data repeated from Figure 4). Dashed line indicates mean MFI (media fluorescence intensity of each sample) value for FL-NPM1c expression after dTAG-13 treatment. **(D)** Heatmap and hierarchical clustering of C-body target genes expression in OCI-AML3 cells with NPM1c overexpression. **(E)** Immunostaining of CD34+ cord blood cells expressing NPM1c with antibodies targeting NUP98 and MENIN. **(F)** Mean CD11b fluorescence level in CD34+ cord blood cells expressing *NPM1c*. **(G)** Representative histogram of CD11b fluorescence intensity of CD34+ cord blood cells expressing FL-NPM1c, C-body null, and mCherry. **(H)** Schematic depicting NPM1c overexpression in murine HSPCs harboring mutant *Dnmt3a.* **(I)** Fold change of donor cells expressing NPM1c in the peripheral blood of mice (*n* = 3 per group). **(J)** Schematic depicting the role of C-bodies in human and murine primary cells. All images are shown in Fire LUT except colocalization (cyan and magenta). White scale bars = 2µm. Error bars = mean ± standard deviation. *p <.05, **p <.01, ***p <.001, Z-test.

Indeed, FL-NPM1c, ΔA3, and ΔB2 form condensates at higher concentrations indicative of a C_sat_, however we did not observe consistent C-body formation at any expression level for the remaining truncations (Figure 6B). Importantly, all proteins had strong enrichment in the cytoplasm, including the C-body null variant (Figure 6A**, S5H**). Together these results indicate that C-body formation depends on several domains of NPM1c, consistent with multivalent interactions needed for heterotypic phase separation.

Having established that some truncations are able to form C-bodies in the absence of endogenous NPM1c, we next asked which variants could prevent cells from differentiating, preserving a key leukemic feature. After expressing all truncated proteins, we degraded endogenous NPM1c and measured a monocytic differentiation marker (CD11b) against a DMSO-treated control after two weeks.

Cells lacking C-bodies – those which expressed the fluorescent protein control mCherry, ΔA2, ΔB1, ΔCTD, and NPM1wt+NES – showed substantial increases in CD11b expression (Figure 6C). Strikingly, cells expressing FL-NPM1c, ΔA3, or ΔB2 showed minimal CD11b expression, maintaining an undifferentiated leukemic state (Figure 6C). Importantly, cells expressing the C-body null variant also differentiated (Figure 6C). The observation that all constructs showed cytoplasmic localization, but only those that contributed to C-bodies were able to prevent differentiation, indicates that C-body formation is required to prevent cells from undergoing differentiation. NPM1c variants that are solely cytoplasmic are insufficient. Together, these data reveal that strong partitioning (K) and condensate formation ability (C_sat_) are predictive of a protein’s ability to prevent differentiation and maintain the leukemic state.

To validate the connection between C-bodies, monocytic differentiation and the leukemic transcriptional program, we analyzed RNAseq data obtained from *NPM1*^WT/Degron^ OCI-AML3 cells expressing FL-NPM1c, “C-body null”, or mCherry constructs after endogenous NPM1c degradation with dTAG-13. We clustered the samples based on the expression of NPM1c-specific genes (**Figure S3B**) revealing that only FL-NPM1c maintains gene expression after NPM1c degradation, in contrast to “C-body null” and mCherry alone (Figure 6D). This establishes the C-bodies as necessary for maintaining the leukemia-associated transcriptional program.

Next, we examined whether *de novo* induction of C-bodies in normal human blood cells would lead to the establishment of the NPM1c-specific expression program and leukemic features. We introduced FL-NPM1c,“C-body null”, or mCherry into progenitors (CD34+) isolated from human umbilical cord blood. As expected, NPM1c, but not the others, formed *de novo* C-bodies evidenced by focal colocalization of NUP98 and MENIN (Figure 6E). When allowed to differentiate toward monocytic lineages in the presence of conditioned media from stromal cells (HS-5), only FL-NPM1c was able to prevent monocytic differentiation (Figure 6F**, G and S6D**). These data underscore the necessity of C-bodies for maintaining a stem-cell-like state and demonstrate that cytoplasmic localization alone is dispensable.

Finally, we sought to determine if C-bodies enable engraftment of NPM1c AML cells and are responsible for their expansion *in vivo.* We overexpressed FL-NPM1c, “C-body null” NPM1c, or mCherry in *Dnmt3a* mutant murine HSPCs and transplanted them into the WT recipient mice (Figure 6H). All genetic variants were able to engraft, however, only cells expressing FL-NPM1c expanded in the peripheral blood between weeks 2 and 4 and persisted in the blood (**Figure 6I and S6E**). “C-body null” and mCherry cells were either depleted or stayed at the same level over 10 weeks.

Thus, we conclude that NPM1c phase-separation is necessary and essential for the maintenance of the stem-like phenotype observed in leukemia cells, *HOX* gene up-regulation, and cellular expansion *in vitro*, *ex vivo* and *in vivo* in both human and murine cells.

Plots depict phase diagrams (based on analytical solutions to Flory-Huggins theory shown in Figure S7) in which co-expression of individual proteins yields co-existing (distinguishable) or mixed (indistinguishable) condensates. Shaded regions indicate distinct phases. Dots indicate NPM1c condensates (red), oncofusion condensates (blue), co-existing condensates (red and blue), or mixed condensates (purple). Points on axes represent cells expressing only one protein and have been shifted (dashed line). **(E)** Live cell imaging of U2OS cells expressing NPM1c and individual oncofusion proteins. **(F-H)** Quantification of nucleoplasm in individual cells expressing oncofusions, NPM1c, or both (Photons @ R.S.). Black dots indicate cells without nuclear condensates, and remaining colors are as indicated in (D). Shaded regions highlight approximate phase boundaries. Points on axes represent cells expressing only one protein and have been shifted (dashed line). NPM1c data from x-axes is duplicated and shown in all 3 panels. All images are shown in Fire LUT except colocalization (cyan and magenta). White scale bars = 2µm. All insets are magnified 2µm square regions.

### Multiple leukemogenic fusion proteins drive C-body formation

Our data has demonstrated that NPM1c forms nuclear C-bodies enriched with multiple proteins that facilitate leukemogenesis. Notably, chromosomal rearrangements of Nucleoporin and *KMT2A* can generate oncofusion proteins implicated in subtypes of leukemias similar to that driven by NPM1c with characteristic *HOXA/MEIS1* gene expression^60–62^. NUP98-fusion proteins have been shown to form condensates in leukemia, and additional studies have reported on the potential of KMT2A oncofusions to phase separate^15,18^. Indeed, nucleoporin and KMT2A fusion proteins are thought to work alongside MENIN and XPO1 to regulate gene expression^55,63^. Given the considerable overlap of protein complexes associated with leukemias driven by NPM1c and these oncofusions, we sought to determine if oncofusion-mediated condensates also coordinated recruitment of XPO1, NUP98, KMT2A, and MENIN and whether these condensates were structurally similar to NPM1c-driven C-bodies.

We first expressed NUP98::NSD1, and SET::NUP214 oncofusions in U2OS cells, and observed *de novo* nuclear condensate formation (Figure 7A) consistent with previous reports^15,63^. Similarly, we expressed the KMT2A::AFF1 oncofusion in U2OS cells and observed previously unreported nuclear condensates (Figure 7A). We next asked whether these oncofusions recruited proteins associated with NPM1c-driven C-bodies. Immunostaining revealed that NUP98::NSD1 and SET::NUP214 condensates were sufficient to recruit XPO1, NUP98, KMT2A, and MENIN (Figure 7B-C**, S7A-B**), consistent with our observations in U2OS cells expressing NPM1c (**Figure S2D-E**). KMT2A::AFF1 condensates were sufficient to recruit XPO1, NUP98, and MENIN, however KMT2A enrichment was not observed. Moreover, we examined LOUCY cells – a T-cell acute lymphoblastic leukemia (T-ALL) line – which harbor nuclear SET::NUP214 condensates. We observed focal enrichment of XPO1, KMT2A, and MENIN consistent with previous reports^63^ and identified NUP98 recruitment to these condensates (**Figure S6C-D**). Importantly, HL-60 cells, which lack nucleoporin or *KMT2A* oncofusion-encoding mutations, did not display highly enriched nuclear puncta (**Figure S7C-D**).

**Figure 7:**
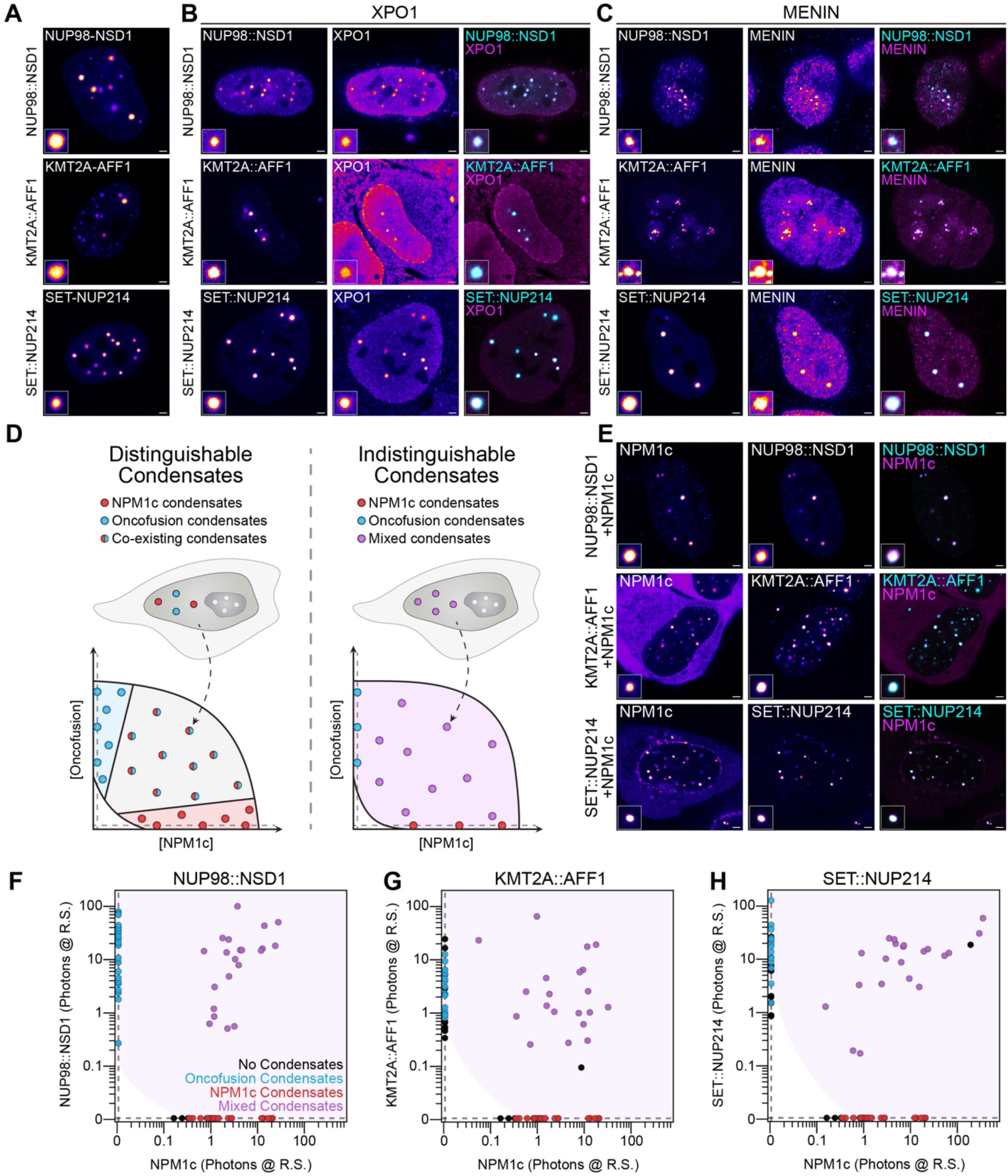
Independent drivers of phase separation converge on C-body formation in leukemia. **(A)** Live cell imaging of U2OS cells expressing oncofusion proteins. **(B)** Immunostaining of U2OS cells expressing oncofusion proteins with antibodies targeting XPO1 and **(C)** MENIN. **(D)** Schematic representation of the biophysical assay to identify condensate miscibility in cells.

Finally, we asked whether these oncofusion condensates are indeed C-bodies. Our observation that NPM1c and oncofusion condensates recruit a common set of proteins strongly supports their convergence on C-bodies. However, separate co-existing condensates can share protein partners, such as DDX6 in P-bodies and stress granules, or Fibrillarin in Cajal bodies and nucleoli^24,64,65^. To determine if multiple condensates are scaffolded from similar interactions in cells, we formalized our observations into a new quantitative miscibility assay, grounded in polymer physics, that experimentally probes the phase diagram for independent drivers of phase separation (**Figure S7E-I**).

For example, if interaction networks driving oncofusion and NPM1c condensates are separable, then two distinct phases (i.e. two types of co-existing condensates) will be achievable within a cell. In contrast, if interaction networks are highly interdependent, then only one phase will ever be achieved (i.e. one type of mixed condensate) within a cell, despite some compositional differences between different cells. To avoid loss of generality under this framework, we define these two cases as biophysically distinguishable or indistinguishable respectively (Figure 7D).

Employing this new assay, we co-expressed oncofusions with NPM1c in U2OS cells, and observed substantial enrichment of oncofusion proteins in NPM1c-driven C-bodies (Figure 7E). At all expression levels where phase separation occurs, we observed a single type of mixed condensate containing NPM1c and oncofusion proteins within each cell, indicating that these condensates are biophysically indistinguishable nuclear structures (Figure 7F-H). Together with the recruitment of a shared network of interaction partners, these findings imply that nucleoporin/KMT2A oncofusion condensates are indeed C-bodies. As demonstrated here in *NPM1*-mutant AML, we propose that C-bodies driven by oncofusions may play a role in driving disease across multiple subtypes of leukemia.

## Discussion

*NPM1* mutations drive approximately one third of newly diagnosed cases of adult AML^27,28^; but how does cytoplasmic NPM1c paradoxically influence gene expression in the nucleus? What is the relationship between NPM1c localization and its role in leukemic transformation? Do other leukemia subtypes with similar gene expression profiles share a mechanistic basis? In this study, we show that NPM1c forms phase-separated nuclear condensates – termed coordinating bodies or C-bodies – across multiple models of *NPM1*-mutant AML. We find that C-bodies are necessary for driving *NPM1*-mutant AML. Finally, we report that leukemogenic oncofusion proteins found in other subtypes of leukemias form nuclear structures which are biophysically indistinguishable from NPM1c-driven C-bodies, suggesting that C-bodies are a unifying feature of multiple subtypes of leukemia.

### C-bodies drive *NPM1*-mutant AML

Despite its pathognomonic cytoplasmic localization, NPM1c’s essential place of action for driving leukemia has remained murky. Indeed, studies over the last two decades have focused on aberrant localization of NPM1c that may lead to loss-of-function of nuclear and nucleolar proteins or gain-of-function in the cytoplasm and nucleoplasm^36–38,55,66–69^. Here, we show for the first time that NPM1c forms nuclear C-bodies, a new biomolecular condensate observed across *in vitro* and *in vivo* models of *NPM1*-mutant AML, and multiple primary *NPM1*-mutant AML patient samples. Our data demonstrate that C-bodies are necessary for all the hallmarks of this leukemia: maintaining *HOXA* gene expression, preventing differentiation, and promoting expansion *in vivo*. Critically, we report that NPM1c localization in the cytoplasm, nucleoplasm, or nucleolus is not sufficient to promote the leukemic phenotype, establishing C-bodies as the primary driver of leukemogenesis.

How do C-bodies work? Previous reports have shown that NPM1c degradation leads to rapid reduction of nascent transcripts from NPM1c-bound target genes and alters chromatin modifications^35–37^ yet a specific mechanism of action has not been resolved. In our previous work, we speculated that NPM1c may act on chromatin^35^. To that end, we identified several proteins – including XPO1, NUP98, KMT2A, and MENIN – that localize to NPM1c-driven C-bodies and *HOXA/MEIS1* chromatin. Indeed, C-bodies are enriched at active chromatin regions including at the *HOXA9* locus. Critically, NPM1c is sufficient for *de novo* C-body formation that recruits associated proteins in multiple genetic contexts *in vitro* and *in vivo*. C-body destabilization or changes in composition prevent the recruitment of these proteins to distinct nuclear structures leading to reduced chromatin binding, lower *HOXA/MEIS1* gene expression, and a shift toward monocytic differentiation. Our extensive truncation series reveals that C-body formation is essential for maintaining the leukemic phenotype and that cytoplasmic NPM1c is not sufficient. Indeed, employing the C-body null variant of NPM1c, we are able to show that C-bodies are essential for cellular expansion *in vitro* and *in vivo*. Together, our data suggest that C-bodies coordinate the recruitment of NUP98 and other proteins such as the COMPASS-like methyltransferase complex – which includes KMT2A and MENIN – to activate developmentally regulated genes in leukemia^60,70^.

Our study reveals new insights into the mechanism of action of targeted therapies with clinical promise in *NPM1*-mutant AML. XPO1-inhibitors break the NPM1c-XPO1 interaction, leading to relocalization of NPM1c from the cytoplasm to the nucleolus as previously described^29,31,71^. Here we further demonstrate that XPO1-inhibition causes C-body dissolution. Our study decouples NPM1c-XPO1 interaction and cytoplasmic export, and suggests that the activity of XPO1-inhibitors in treating disease is a result of C-body disruption rather than export inhibition. Similarly, MENIN inhibitors are a leading class of therapies that break the KMT2A-MENIN interaction and disrupt aberrant chromatin activation, yet their precise mechanism in *NPM1*-mutant AML is not well defined^40–42,56^. We show that MENIN inhibitors cause loss of MENIN - but not KMT2A - from C-bodies, despite the persistence of MENIN puncta elsewhere in the nucleus. The effectiveness of MENIN inhibitors in downregulating *HOXA* gene expression and promoting differentiation indicates that the critical interaction between MENIN and KMT2A occurs within C-bodies. MENIN inhibitor treatment reduces expression of C-body bound genes and shifts cells toward monocytic differentiation in *in vitro* and *in vivo* models and primary *NPM1*-mutant AML patient samples. Broadly, our results indicate that therapeutic targeting of C-body formation or function may be a promising strategy in *NPM1*-mutant AML.

Our findings have implications for additional aspects of *NPM1* mutation in disease. Notably, heterozygous loss of *NPM1* (e.g. del(5q)) is associated with myelodysplastic syndromes and genome instability, yet *NPM1*-mutant AMLs typically have a normal karyotype^72^. Our observation that NPM1c colocalizes with NPM1wt at the chromosomal sheath during mitosis may indicate that its function in maintaining genome stability is preserved. Separately, our observation that NPM1c overexpression drives *de novo* C-body formation suggests that co-mutations normally found in *NPM1*-mutant AML (e.g. in *DNMT3A* or *TET2*)^73,74^ are not required for C-body formation but instead functionalize C-bodies through altered gene expression or chromatin accessibility.

### Heterotypic Interactions Dominate NPM1 Phase Separation

Phase separation driven by multivalent weak interactions is a general paradigm underlying many membrane-less compartments within cells^23,46^. In this context, our observation of the strong spatial separation between NPM1c and NPM1wt is largely unexpected, as these proteins ostensibly share weak interactions. Strikingly their separation is imposed by the cell cycle where during telophase, NPM1c and NPM1wt relocate from the chromosomal sheath to C-bodies and PNBs, respectively. Throughout interphase, this separation is enforced by the immiscibility (i.e. exclusivity) of distinct interaction networks (e.g. rRNA in ribosomal intermediates or the NUP98/COMPASS complex) coordinated by NPM1wt and NPM1c.

How does a small change in amino acids at the C-terminus encode this separation? NPM1 has a self-pentamerization domain which enables multivalent interactions through its charge blocky IDR and RBD domains. The IDR, unchanged in NPM1c, facilitates recruitment to the chromosomal sheath during mitosis^75^, consistent with our observations. In contrast, the RBD is essential for NPM1wt’s role in nucleolar formation, and is mutated into an XPO1-binding NES in NPM1c^22,26,27^. While our data suggests that interaction with XPO1 is sufficient to decimate enrichment into nucleoli, this interaction is insufficient to form *de novo* C-bodies. Indeed our truncation series illuminates multiple regions of NPM1c necessary for C-body formation. This suggests that additional interaction partners, which have less affinity for NPM1wt, contribute to C-body formation.

Endogenous condensates rely on interaction networks (a.k.a. heterotypic interactions) and thus exhibit composition-dependent stability^22,24^. Similarly, we find that overexpression (i.e. compositional overrepresentation) of NPM1c and relevant truncations destabilizes C-bodies and results in decreased cell growth. Together, these data demonstrate that C-bodies are also driven by heterotypic interactions.

### C-bodies as a Common Therapeutic Target Across Leukemia Subtypes

Phase separation is an emerging feature of many leukemias, where dozens of genetic alterations involving nucleoporin genes, *KMT2A*, *ENL* and others are reported to cause condensate formation^14,15,18,60,61,76,77^. Broadly, hundreds of mutant proteins are proposed to drive phase separation in disease, but whether there is functional redundancy between condensates remains an open question^17^. Consolidating unrelated mutant proteins as independent drivers of the same functional condensate would clarify disease mechanisms. Toward this end, we develop a simple assay - grounded in polymer physics - to determine whether independent proteins form the same indistinguishable condensate (**Figure S7E-I**). Remarkably, this approach revealed that many leukemic oncofusions also drive C-body formation, suggesting that C-bodies are a common feature of many leukemias. We speculate that phase separation underlies aberrant chromatin regulation in a growing list of leukemias driven by fusion oncoproteins^78^. We anticipate that this conceptual framework will be applicable in many contexts and enable the identification of pathogenic condensates underlying multiple diseases.

How do the principles underlying multi-component phase separation inform strategies for targeting oncogenic condensates? We show that C-bodies rely on a network of interactions to form and function. We anticipate that new or existing drugs disrupting proteins in this network will destabilize condensates and therefore demonstrate clinical activity. As an example, XPO1 and MENIN inhibitors have already shown activity across *NPM1*-mutant, *NUP98*-fusion, and *KMT2A*-rearranged leukemias, likely owing to their shared impact on C-bodies^40–42,52,71,79^. Notably, some patients develop mutations at the MENIN-KMT2A interface that confer resistance to MENIN-inhibitors, highlighting the value of new approaches targeting C-bodies^80^. Future investigation into the composition and dynamics of C-bodies and other pathogenic condensates will be crucial for the development of effective therapies against disease.

## Acknowledgements

We thank Jason Lee for providing microscopy access for pilot studies, the Leukemia Sample Bank at the University of Texas MD Anderson Cancer Center for access to primary patient samples, and Liling Wan and Kyle Eagen for help with DNA FISH experiments. We also thank all members of the Riback and Goodell laboratories for useful comments and feedback. This project was supported by CA183252 (M.A.G.), CA26574 (M.A.G.) and F30CA268725 (G.K.D.). J.A.R. is a CPRIT Scholar in Cancer Research with funding from the Cancer Prevention and Research Institute of Texas (CPRIT) New Investigator Grant RR210040 and was supported by The Ted Nash Long Life Foundation, the Leukemia Research Foundation, and the Searle Scholars Program. R.E.R. was supported by K08CA201611 and K12CA090433-11. B.F. was supported by Associazione Italiana per la Ricerca sul Cancro (AIRC). L.B. was supported by Associazione Italiana per la Ricerca sul Cancro (AIRC) Start-up Grant nr.22895. VA was supported by the Trond Mohn Foundation and The Research Council of Norway (project no. 303358). BTG was supported by the Norwegian Cancer Society (grants numbers 303445 and 19017), The Research Council of Norway (project no. 303358) and Helse Vest Health Trust (project no. F-13102). This work was also supported by the Cytometry and Cell Sorting Core with funding from CPRIT and the NIH (RP180672; CA125123, RR024574). N.S. was supported by the Andrew Sabin Family Foundation Fellowship. S.S.Y. was supported by a Scialog Award sponsored jointly by the Research Corporation for Science Advancement and the Gordon and Betty Moore Foundation (award number 28418). N.S. is a CPRIT Scholar in Cancer Research with funding from the Cancer Prevention and Research Institute of Texas (CPRIT) New Investigator Grant RR160021, and RP220292.

## Author Contributions

Conceptualization, G.K.D., M.A.G. and J.A.R.; Methodology, G.K.D., A.A., E.K., C.-W.C., E.P., M.A.G. and J.A.R.; Investigation, G.K.D., A.A., E.K., J.B., M.S., N.C., G.L., C.D., A.M., A.G., K.W.; N.S. Formal analysis G.K.D., E.L., C.-W.C., J.d.l.F, J.A.R.; Data Curation, G.K.D., C.-W.C., S.S.Y., J.d.l.F; N,S, J.A.R.; Resources, A. Maiti, S.S.Y., N.J.S., N.S., V.A., B.T.G., B.F.; Writing - Original Draft, G.K.D., M.A.G., J.A.R.; Writing - Review & Editing, G.K.D., A.A., E.K., J.B., M.S., C.-W.C., E.P., R.E.R., L.B., N.S; M.A.G., J.A.R.; Supervision, R.E.R., L.B., M.A.G., J.A.R.

## Supplemental Data

**Figure S1:**
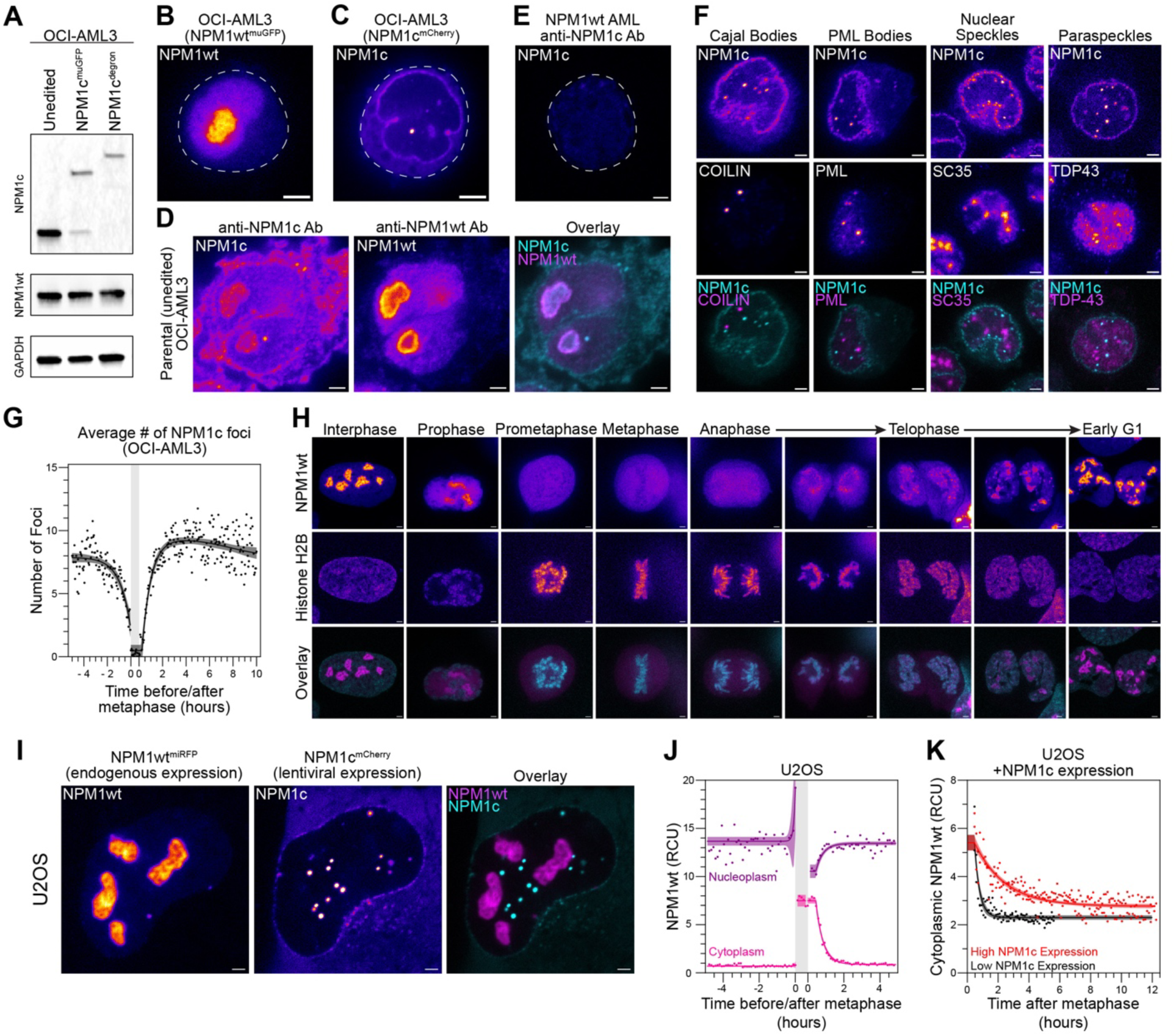
NPM1c forms distinct biomolecular condensates and impairs NPM1wt transit to the nucleus following cell division. **(A)** Immunoblotting of protein lysates from OCI-AML3 cell lines with no editing (Unedited), NPM1c tagging with muGFP (NPM1c^muGFP^) or NPM1c tagged with an inducible degrader tag (NPM1c^degron^) with anti-NPM1c and anti-NPM1wt antibodies. GAPDH is used as an indicator of loading control. **(B, C)** Live cell imaging of CRISPR-edited OCI-AML3 cells containing endogenous C-terminal-tagged NPM1c and NPM1wt (mCherry and muGFP respectively). **(D)** Immunostaining of parental OCI-AML3 cells **(E)** Immunostaining of *NPM1^wt/wt^* mononuclear cells derived from an AML patient without *NPM1* mutation using antibodies targeting NPM1c. **(F)** Immunostaining of *NPM1*^WT/Degron^ OCI-AML3 cells expressing fluorescently labeled NPM1c with antibodies targeting Cajal bodies (COILIN), PML bodies (PML), Nuclear Speckles (SC35), and Paraspeckles (TDP43). **(G)** Quantification of average number of NPM1c foci from live-cell imaging data of dual-labeled OCI-AML3 cells imaged every 3 minutes for ∼10hrs (*n* = 10, same dataset as in Figure 1G-J). **(H)** Live cell imaging of dual-labeled U2OS cells expressing endogenously labeled NPM1wt (mCherry) and Histone H2B (muGFP). Representative images of cell cycle stages shown here. **(I)** Live cell imaging of U2OS cells expressing endogenously labeled NPM1wt (miRFP) transduced with NPM1c (mCherry). **(J)** Quantification of U2OS cells from panel **H** imaged every 7 minutes for ∼10hrs (*n* = 5). **(K)** Quantification of U2OS cells from panel **I** measuring NPM1wt (miRFP) in the cytoplasm up to 12 hours after the end of metaphase. Cells with high NPM1c expression are annotated in red, while cells with low NPM1c concentration are in black. Metaphase (shaded gray) is defined by the average time between the earliest visualization of prometaphase and anaphase. Timepoints reflecting prometaphase through anaphase have been annotated as cytoplasm. All concentrations measured by photons in relative concentration units (RCU). All images are shown in Fire LUT except colocalization (cyan and magenta). White scale bars = 2µm.

**Figure S2:**
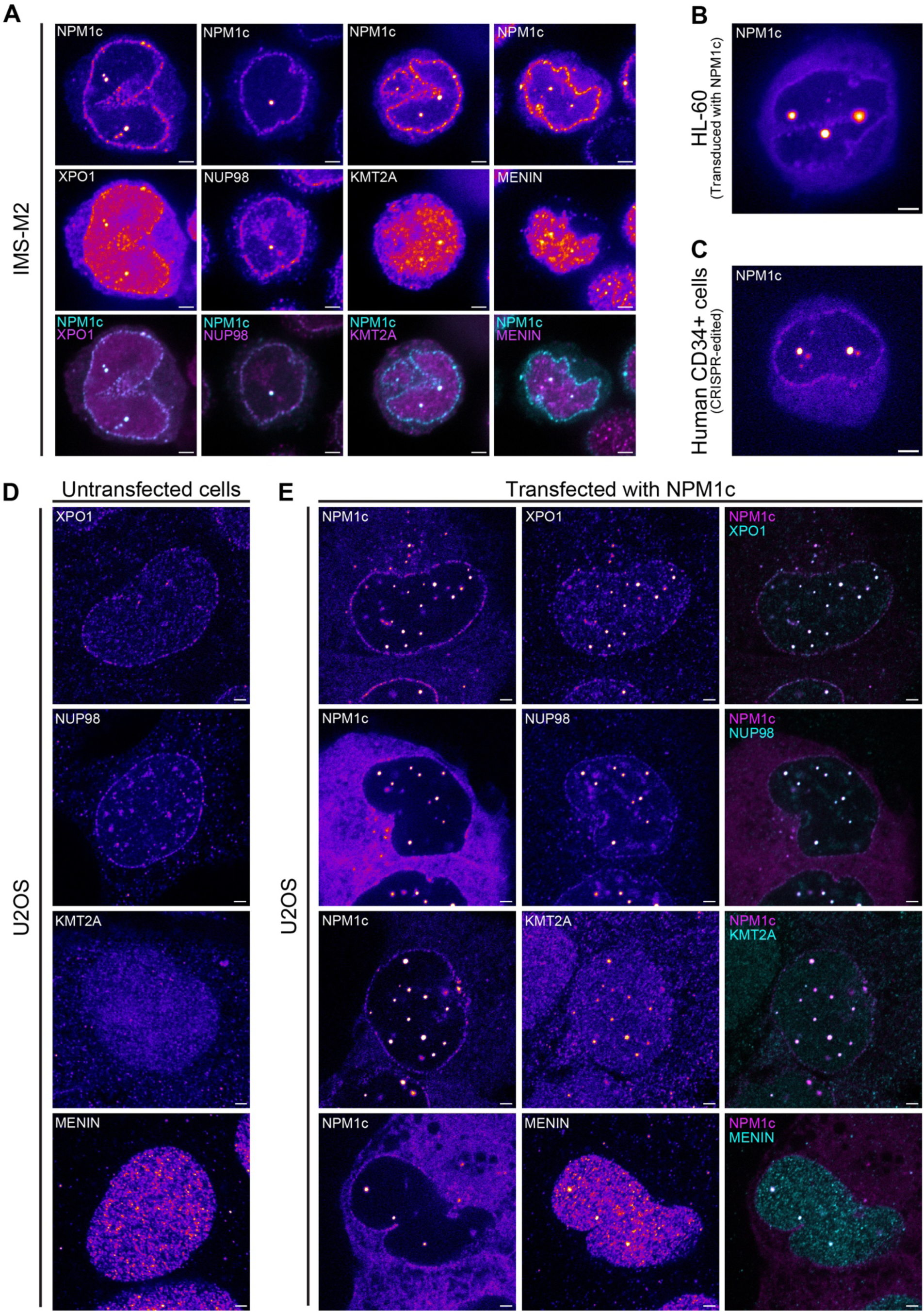
NPM1c is necessary and sufficient for nuclear condensate formation and punctate enrichment of C-body associated proteins. **(A)** Immunostaining of *NPM1*^WT/Degron^ IMS-M2 cells expressing fluorescently labeled NPM1c with antibodies targeting XPO1, NUP98, KMT2A, and MENIN. **(B)** Live cell imaging of parental HL-60 cells after viral transduction with FL-NPM1c. **(C)** Live cell imaging of human CD34+ cells derived from cord blood after CRISPR-editing of NPM1wt to NPM1c. (**D)** Immunostaining of parental U2OS cells and (**E**) parental U2OS cells transfected with NPM1c with antibodies targeting XPO1, NUP98, KMT2A, and MENIN. All images are shown in Fire LUT except colocalization (cyan and magenta). White scale bars = 2µm.

**Figure S3:**
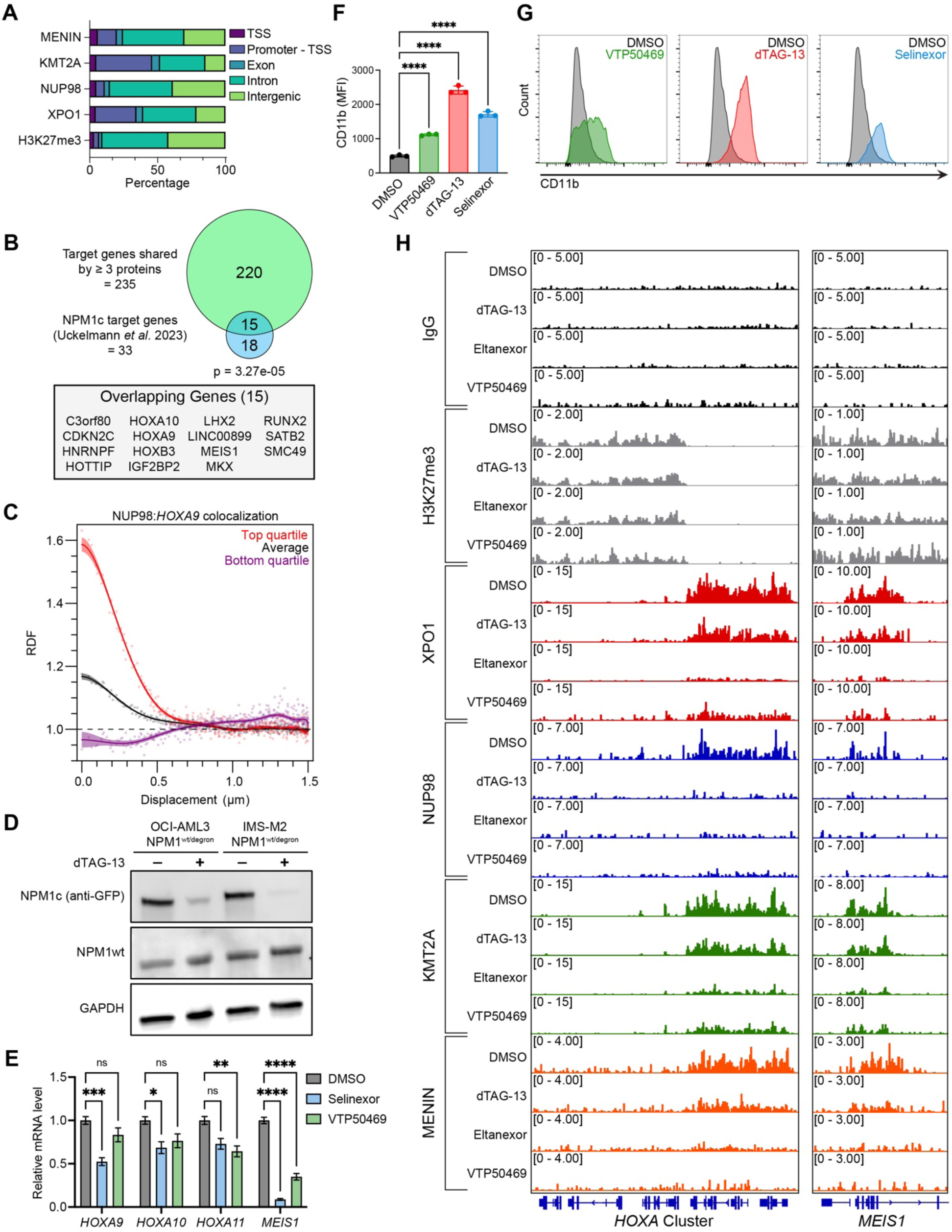
Condensates destabilization is associated with decreased chromatin-binding of C-body associated proteins at key genetic loci. **(A)** Stacked bar plots showing distributions of CUT&RUN data from DMSO-treated *NPM1*^WT/Degron^ OCI-AML3 cells, with genomic peaks annotated as TSSs, promoters, exons, introns, and intergenic regions. **(B)** Venn diagram of TSS-promoter peaks from DMSO-treated CUT&RUN sequencing depicting unique genes bound by at least 3 of 4 antibody targets (including XPO1, NUP98, KMT2A, and MENIN) (*n* = 235) and NPM1c target genes identified by ChIP-seq (*n* = 33, Uckelmann *et al.* 2023) and annotated table of genes found in both datasets. **(C)** RDF of NUP98 and HOXA9 in top (red) and bottom (purple) quartiles in comparison to the average (*n* = 69). **(D)** Immunoblotting analysis of protein lysates from OCI-AML3 and IMS-m2 cells treated with DMSO or dTAG for 72h using anti-GFP, anti-NPM1wt and anti-Gapdh-Rhodamine antibodies. **(E)** Relative mRNA levels of *HOXA/MEIS1* in OCI-AML3 cells treated with VTP50469 or Selinexor. **(F)** Mean CD11b level in OCI-AML3 cells treated with VTP50469, dTAG-13, Selinexor or DMSO for 4 weeks and **(G)** representative histograms of CD11b level. **(H)** Representative Integrated Genome Viewer (IGV) tracks (duplicates per sample) for IgG (black), H3K27me3 (Gray), XPO1 (Red), NUP98 (Blue), KMT2A (Green), and MENIN (Orange) treated with DMSO, dTAG-13 (500nM), Eltanexor (100nM), or VTP50469 (300nM) for 24 hours. Fisher’s Exact test was used for statistical testing.

**Figure S4:**
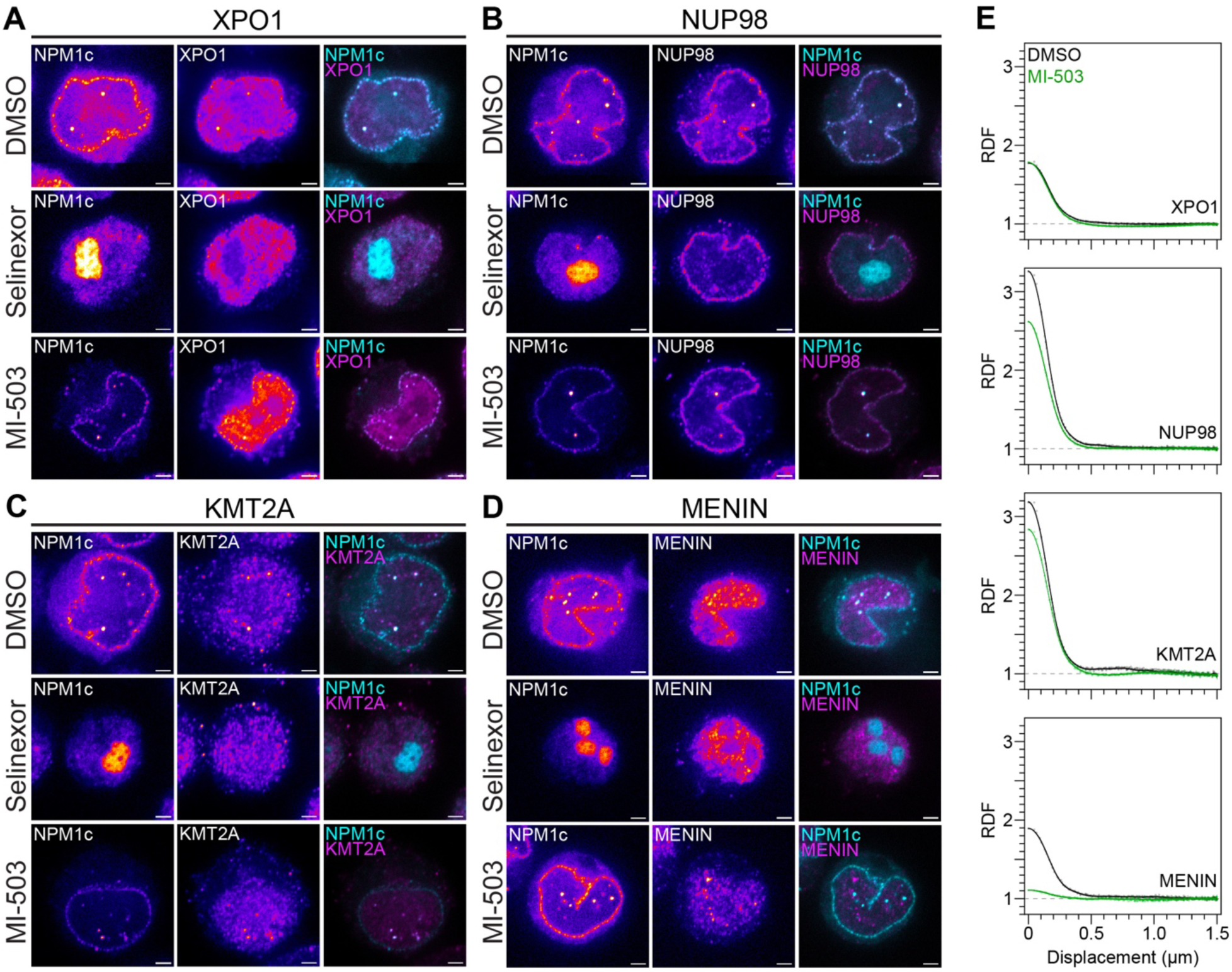
XPO1 and MENIN inhibition disrupt NPM1c condensate formation and composition. **(A-D)** Immunostaining of *NPM1*^WT/Degron^ OCI-AML3 cells with antibodies targeting XPO1, NUP98, KMT2A, and MENIN after DMSO, Selinexor, or MI-503 treatments. (**E)** RDF of XPO1, NUP98, KMT2A, and MENIN with DMSO or MI-503 treatment (*n* = 50). All cells were treated for 24hrs prior to immunostaining (Selinexor = 100nM, MI-503 = 2.5µM). All images are shown in Fire LUT except colocalization (cyan and magenta). White scale bars = 2µm. RDF data for DMSO samples is repeated from Figure 3B.

**Figure S5:**
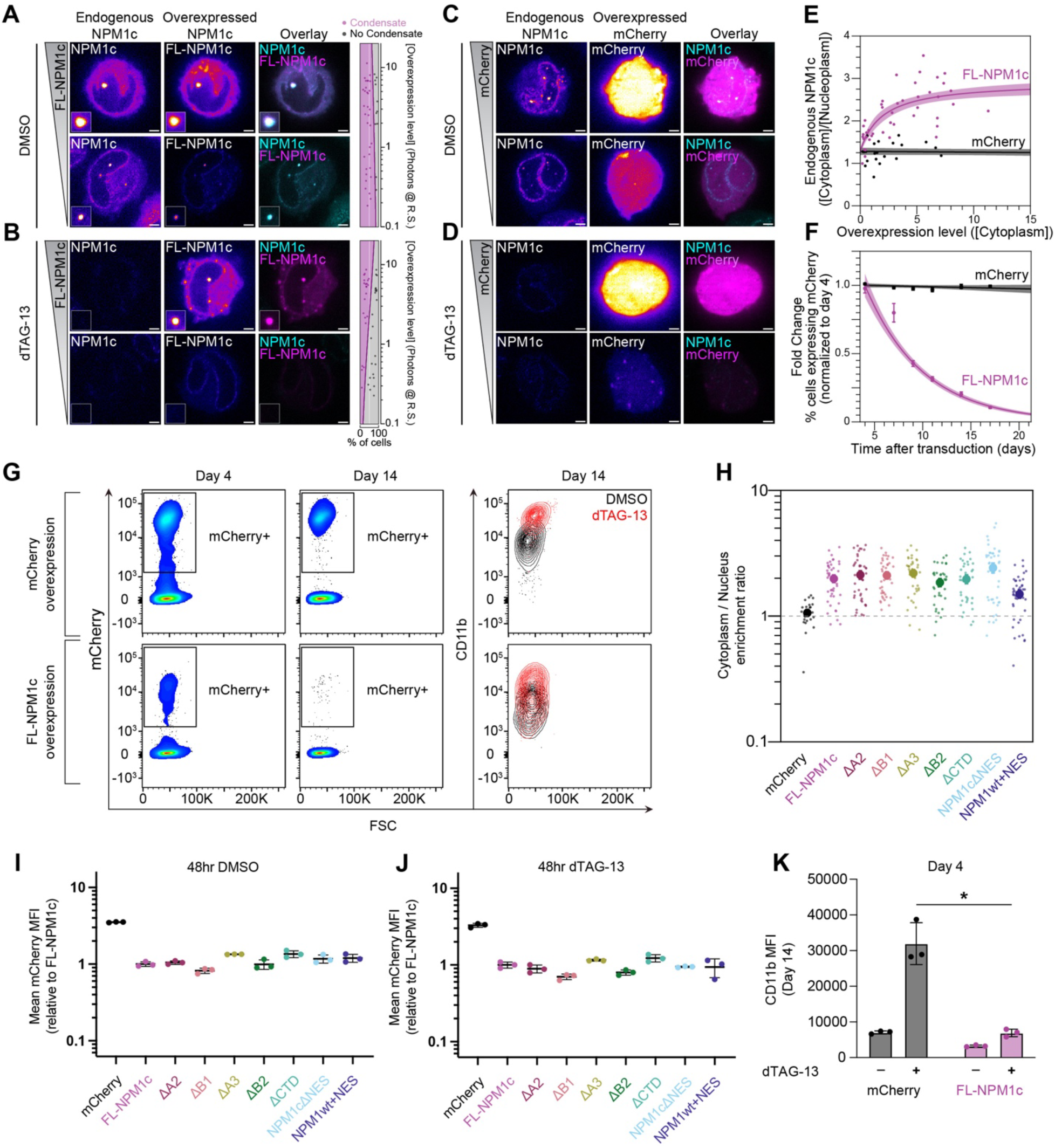
Overexpression of truncated NPM1c proteins and their impact on cell growth and differentiation. **(A)** Live cell imaging of *NPM1*^WT/Degron^ OCI-AML3 cells with stable over-expression of FL-NPM1c treated with DMSO for 24hrs. **(TOP)** high FL-NPM1c expression **(BOTTOM)** low FL-NPM1c expression. **(B)** live cell imaging of *NPM1*^WT/Degron^ OCI-AML3 cells treated with dTAG-13 (500nM) for 24hrs. **(TOP)** high FL-NPM1c overexpression **(BOTTOM)** low FL-NPM1c expression. **(C)** Live cell imaging of *NPM1*^WT/Degron^ OCI-AML3 cells with stable over-expression of mCherry treated with DMSO for 24hrs. **(TOP)** high mCherry expression **(BOTTOM)** low mCherry expression. **(D)** live cell imaging of *NPM1*^WT/Degron^ OCI-AML3 cells treated with dTAG-13 (500nM) for 24hrs. **(TOP)** high mCherry expression **(BOTTOM)** low mCherry expression. **(E)** Correlation of NPM1c and mCherry control overexpression and endogenous NPM1c localization ([Cytoplasm/Nucleoplasm]). **(F)** Normalized fraction of cells with NPM1c or mCherry overexpression over 17 days as measured by flow cytometry. **(G)** Representative flow cytometry plots depicting gating strategy for mCherry+ and CD11b+ fluorescence **(TOP)** Cells expressing mCherry or **(Bottom)** FL-NPM1c were analyzed at days 4 and 14 after viral transduction. **(H)** Quantification of cytoplasm to nucleoplasm ratio of all NPM1c truncations. Dashed line represents a ratio of 1.00. **(I)** Median Fluorescence Intensity (MFI) of mCherry fluorescence via flow cytometry after 48 hours of DMSO or **(J)** 500nM dTAG-13 treatment, *n* = 3 biological replicates per truncation. **(K)** MFI of CD11b via flow cytometry after 14 days of DMSO or 500nM dTAG-13 treatment, *n* = 3 biological replicates per treatment and truncation. (data repeated from Figure 4). *p < 0.05, t test with Welch’s correction. All images are shown in Fire LUT except colocalization (cyan and magenta). White scale bars = 2μm.

**Figure S6:**
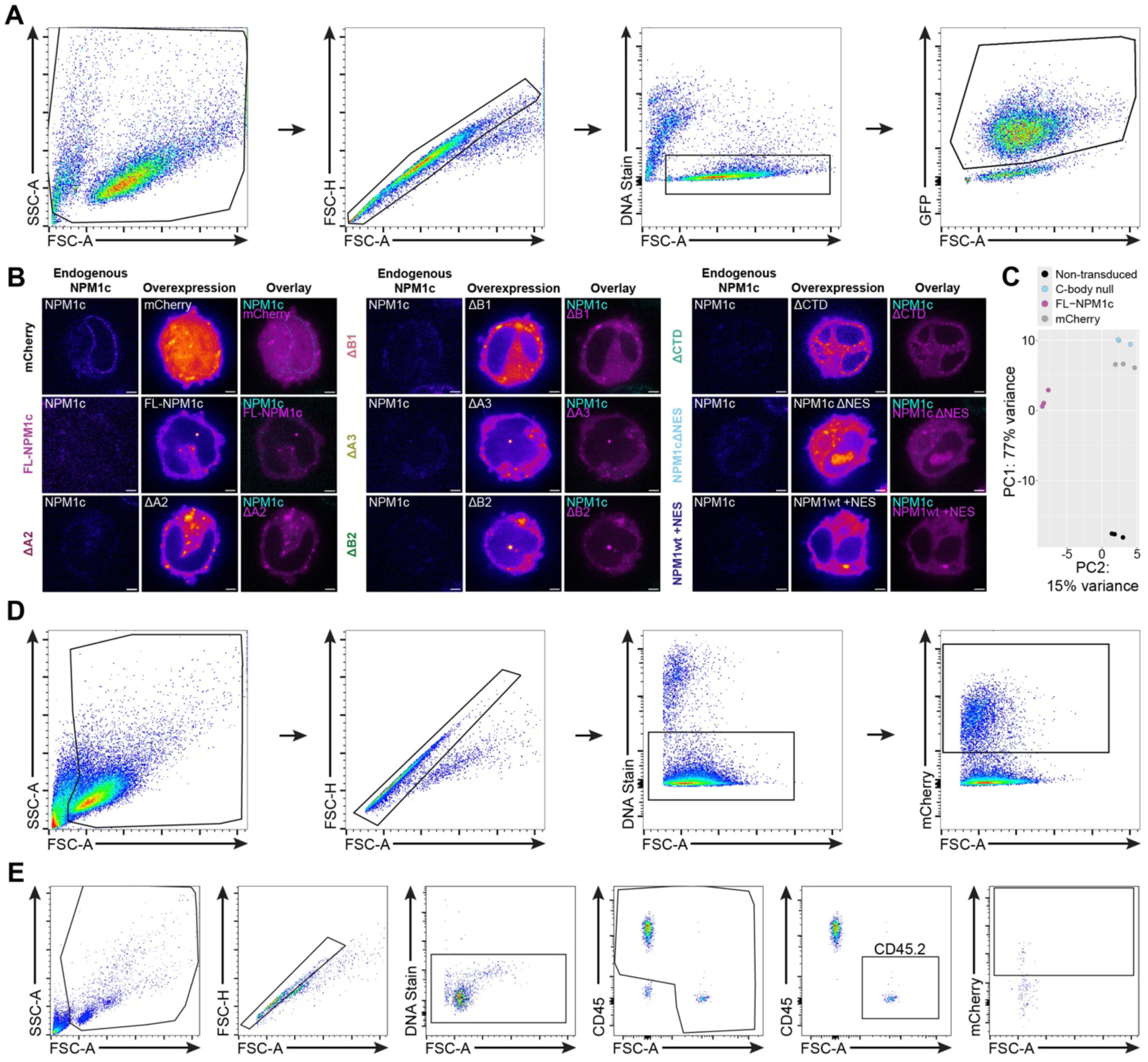
C-bodies in human and murine models of NPM1c AML. **(A)** Gating strategy for murine *Dnmt3a^R878H/+^*;*Npm1^cA/+^*;*Nras^G12D-GFP^*cells **(B)** Live cell imaging of *NPM1*^WT/Degron^ OCI-AML3 cells expressing NPM1c truncations treated with 500nM dTAG-13 for 7 days. **(C)** Principal component analysis (PCA) based on variance-stabilized transcript counts of non-transduced, C-body null, FL-NPM1c, and mCherry samples (*n* = 3). **(D)** Gating strategy for human CD34+ cord blood cells expressing mCherry, FL-NPM1c, or C-body null proteins. **(E)** Gating strategy for murine *Dnmt3a^R878H/+^* cells expressing mCherry, FL-NPM1c, or C-body null proteins. All images are shown in Fire LUT except colocalization (cyan and magenta). White scale bars = 2µm.

**Figure S7:**
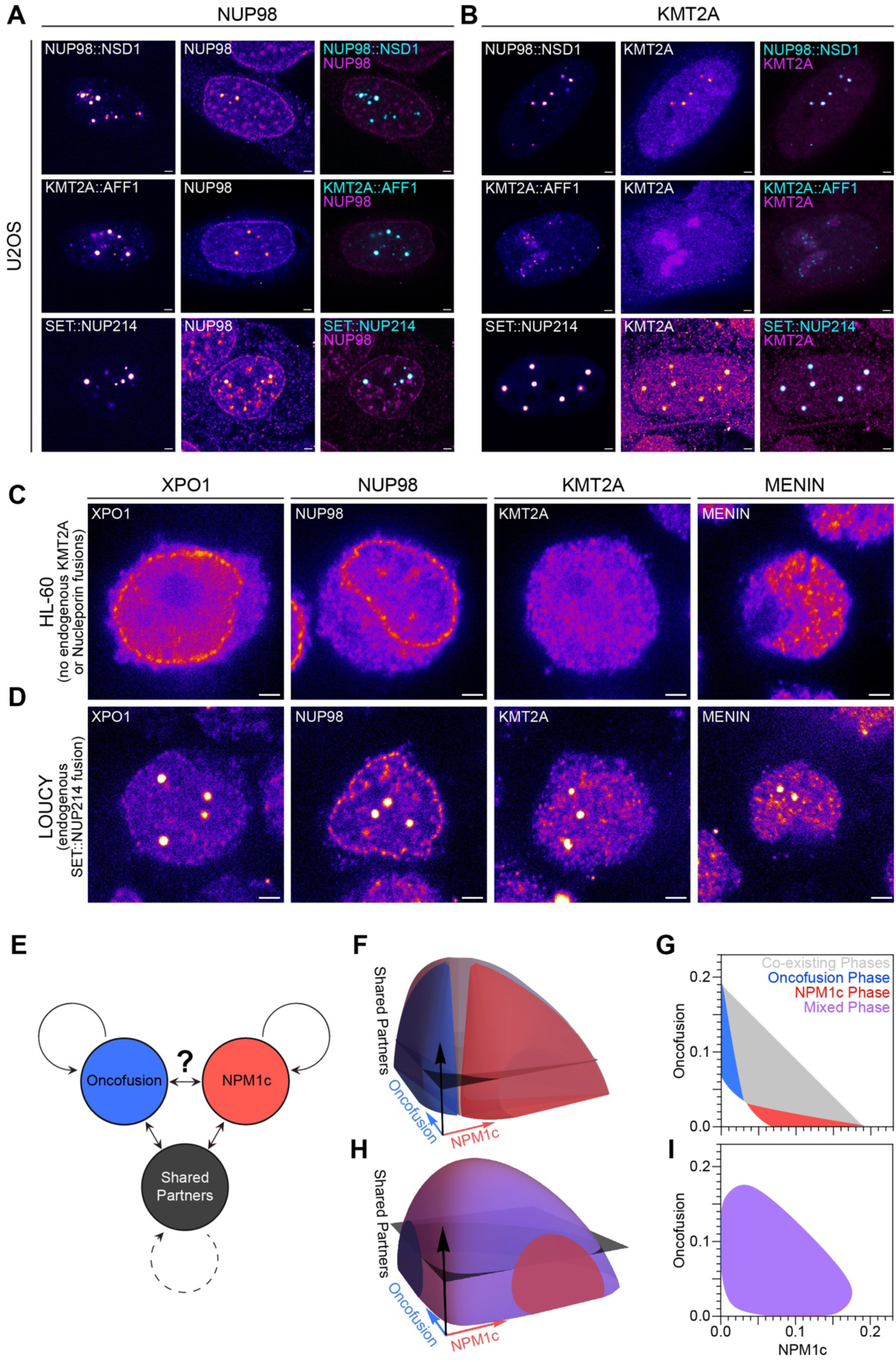
Oncofusion condensates recruit C-body associated proteins and *in silico* validation of the miscibility assay (A) Immunostaining of U2OS cells expressing oncofusion proteins with antibodies targeting NUP98 and **(B)** KMT2A. **(C)** Immunostaining of parental HL-60 cells and **(D)** LOUCY cells with antibodies targeting XPO1, NUP98, KMT2A, and MENIN. **(E)** Schematic representation of 3 components and their interactions. Solid and dashed curved arrows represent a strong (χ = 0.655) and weak (χ = 0.545) tendency to undergo phase separation respectively. Double-headed arrows indicate a favorable interaction (χ = -0.143). Interaction strength between blue and red components (shown as “?”) is modeled as favorable (χ = -0.143) or unfavorable (χ = 0.545) in panels B-C and D-E, respectively. **(F)** Three-dimensional and **(G)** two-dimensional phase diagrams depicting unfavorable interactions between NPM1c and oncofusion components that result in distinguishable phases. **(H)** Three-dimensional and **(I)** two-dimensional phase diagrams depicting favorable interactions between NPM1c and oncofusion components that result in indistinguishable phases. All images are shown in Fire LUT except colocalization (cyan and magenta). White scale bars = 2µm.

**Table S1:**
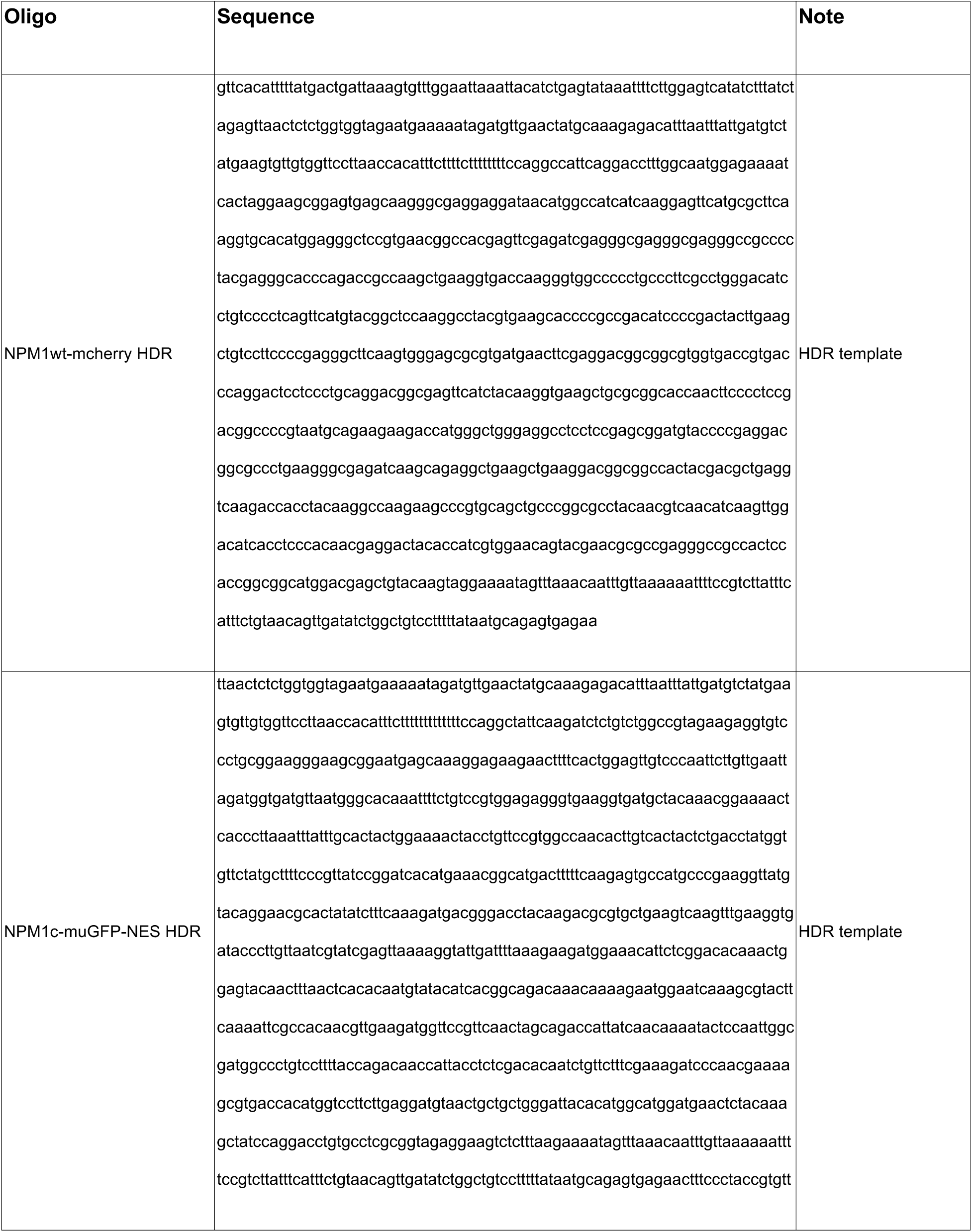
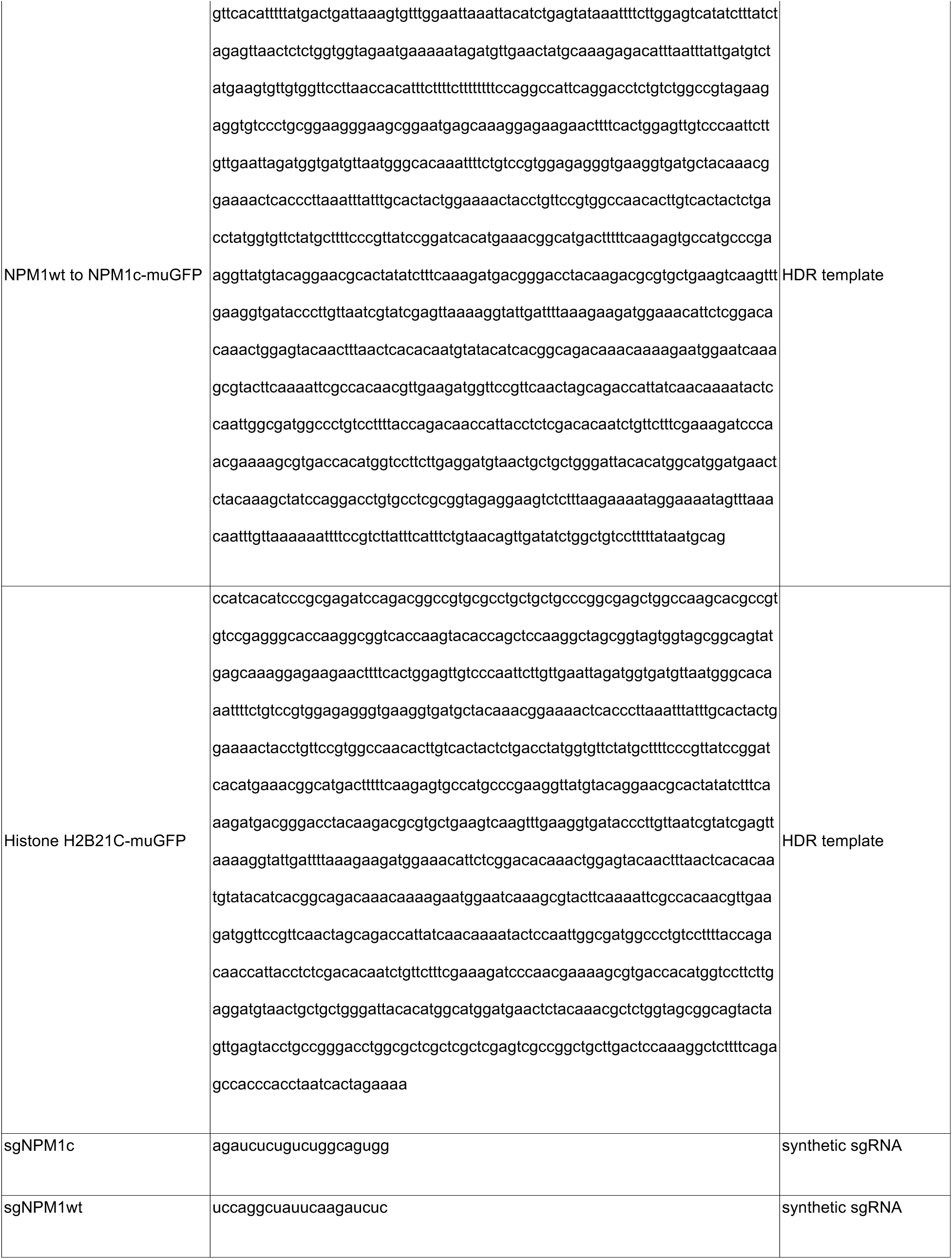
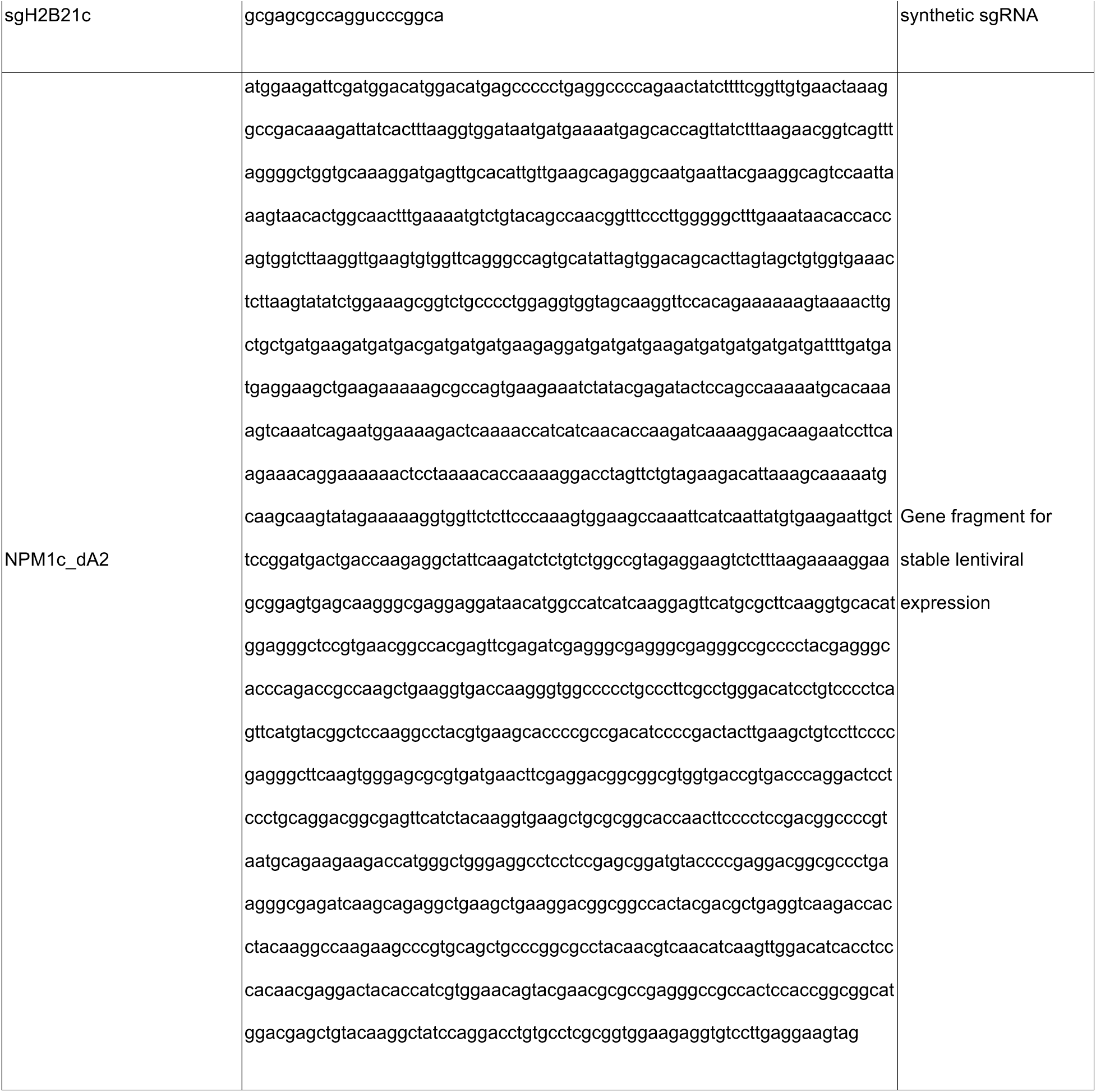
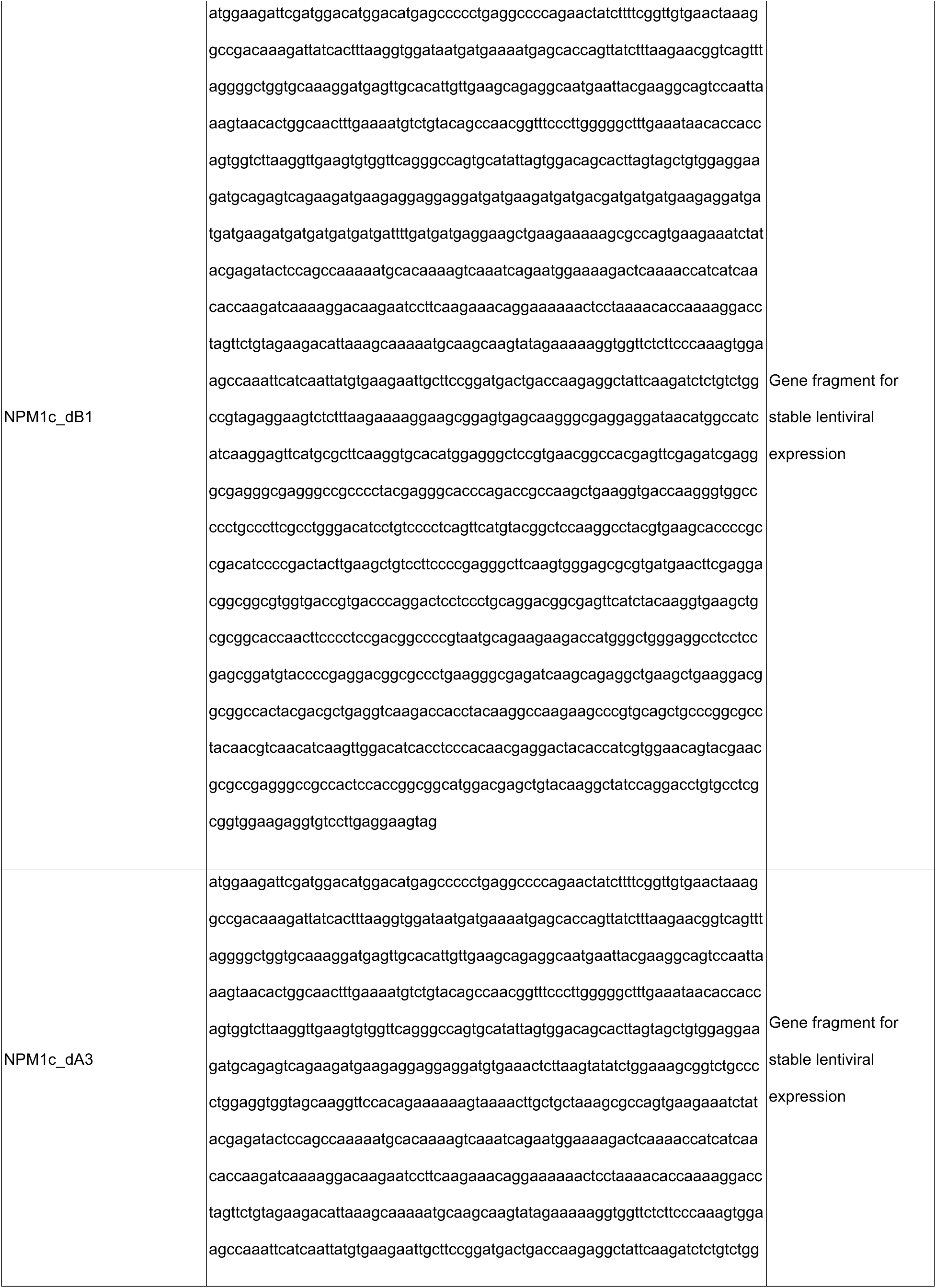
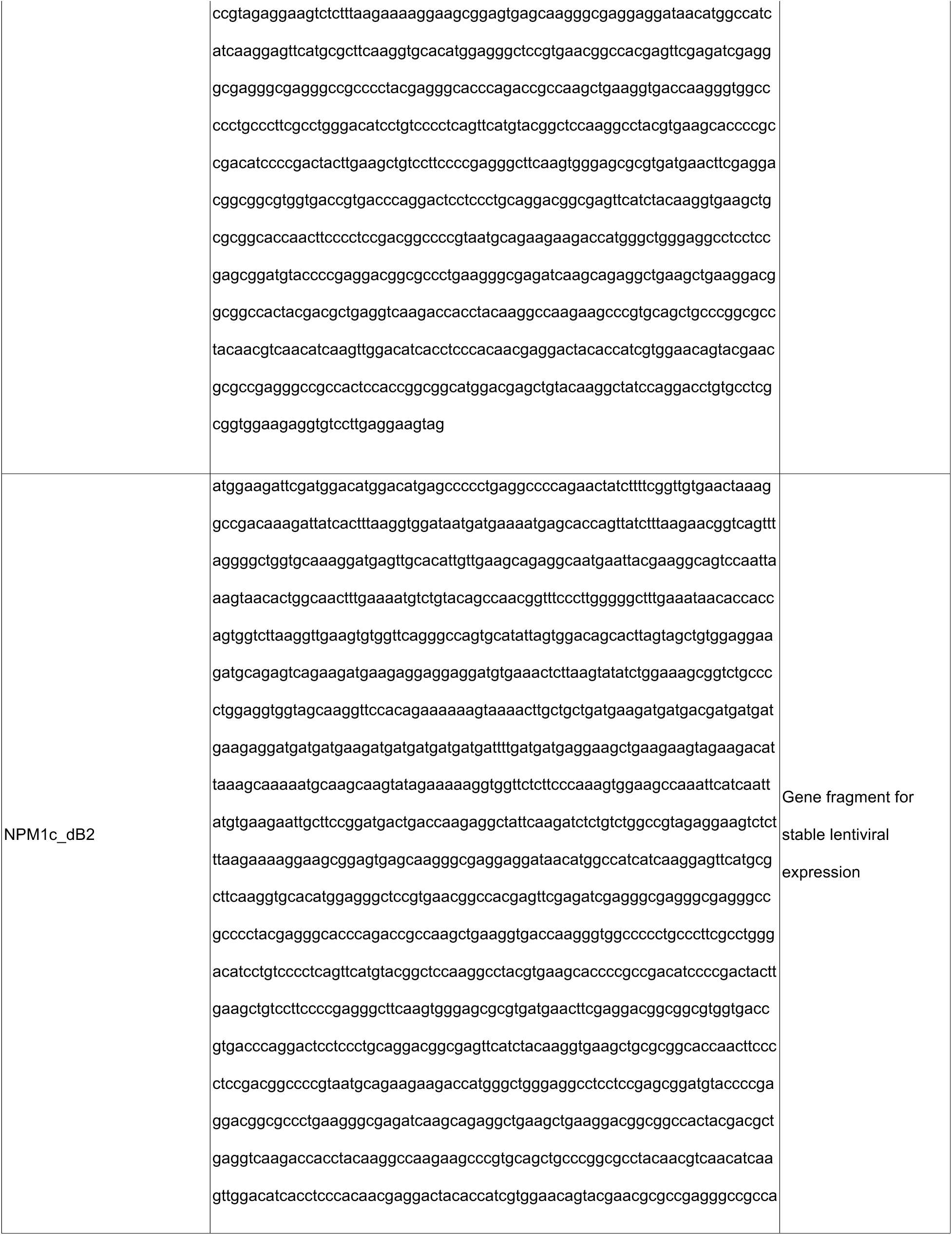
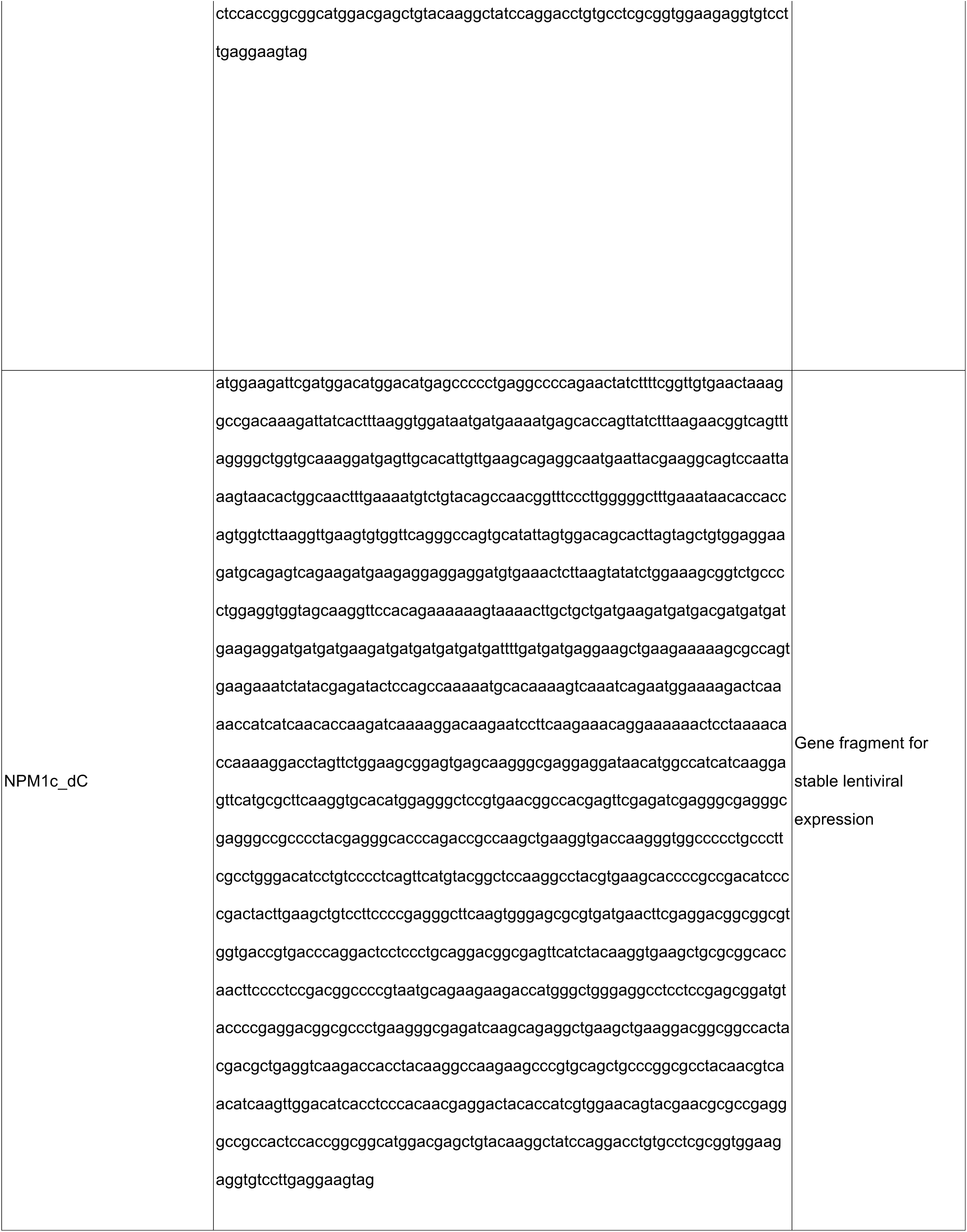
Sequences of custom DNA/RNA oligonucleotides and HDR templates.

**Table S2:** Genomic features of primary AML patient samples (related to Figure S1)

| **Patient number** | **Karyotype** | **Additional Mutations** |
| --- | --- | --- |
| NPM1mut<br>01 | 46,XY,t(8;22)(q24.1;q11.2)[6]/46,idem,+1<br>,der(1;15)(q10;q10)[14] | GNAS R201H;IDH2 R140Q;KRAS<br>G12A;MPL Y591D;NPM1<br>W288fs*12;NRAS;SRSF2P95_R102del |
| NPM1mut<br>02 | PENDING | ASXL1 S1146L;DNMT3A R882C;FTL3<br>F612L + FLT3-ID;IDH2 R140Q;NPM1<br>W288fs;STAT3 D570N ;WT1 D469N |
| NPM1mut<br>03 | 47,XY,+4[20] | DNMT3A;KIT;NPM1;SUZ12;TET2 |
| NPM1wt<br>AML | 46,XX,t(8;21)(q22;q22)[20] | t(8;21) |

## Methods

### Experimental Model and Study Participant Details

#### Cell Culture

All human suspension cell lines (OCI-AML3, IMS-M2, HL-60, and LOUCY cells) were cultured in RPMI 1640 medium supplemented with 10% FBS, 1% penicillin/streptomycin (Pen/Strep). U2OS cells were cultured in DMEM supplemented with 10% FBS and 1% Pen/Strep. Patient Derived Xenograft 2 (PDX2) cells and primary CD34+ cells derived from whole human cord blood were cultured in RPMI media supplemented with 10 mM L-glutamine, 10% FBS, conditioned media from human stromal HS-5 cells. To isolate primary CD34+ cells, mononuclear cells from fresh cord blood samples were isolated using Lymphoprep density gradient (Stemcell technologies). Mononuclear cells were stained with CD34+ beads (130-100-453, Miltenyi Biotech) and purified on MACS columns (Miltenyi Biotech). CD34+ HSCs were frozen in liquid nitrogen for future use.

All cells were cultured in a humidified incubator containing 5% CO_2_ at 37°C. Suspension lines were grown in T-25 flasks and passaged at a 1:6 dilution approximately every 3 days. U2OS cells were grown in 10 cm^2^ dishes and passaged at a 1:10 dilution every 3-4 days with trypsin (Invitrogen, 25300054). PDX2 and primary human CD34+ cells were not passaged, but instead stimulated with 100 ng/µL IL-3 and TPO, 50 ng/µL FLT3-L, 1 µM SR1 (Selleckhem), and 1 µM UM729 (Stemcell technologies) for 48 hours prior to CRISPR/Cas9 experiments.

#### Propagation of Patient Derived Xenograft Model (PDX2)

*NPM1/FLT3*-mutant PDX2 cells were cultured and propagated in NSG mice as described previously ^35^. Briefly, PDX2 cells were thawed and washed twice with PBS. 5x10^6^ cells were injected into irradiated (250cGy) NSG-S (JAX #013062) female mice via tail vein injection. After 3 weeks, mice were sacrificed, and bone marrow cells were stained with human anti-CD45 antibodies (APC, BD Biosciences #561864), and engraftment rate was assessed by flow cytometry. Cells were frozen in liquid nitrogen for future use.

#### Primary AML Samples

Deidentified fresh peripheral blood samples from AML patients with blast count exceeding 60% were purchased from MD Anderson Cancer Center (Leukemia sample bank, MDACC, Houston, TX). The institutional review board approved the procedure and all patients signed a written informed consent before sample acquisition. Sequencing results of a custom panel of 81 genes recurrently mutated in hematologic malignancies, as well as standard karyotyping results were provided by MDACC. Mononuclear cells were isolated using Lymphoprep density gradient (Stemcell technologies) and frozen in liquid nitrogen for future use. Cells were cultured in RPMI media supplemented with 10 mM L-glutamine, 10% FBS and conditioned media from human stromal HS-5 cells. All cells were supplemented with denarase (c-LEcta) for 1 hour after thawing for all experiments.

### CRISPR/Cas9 Genetic Engineering

#### Generation of Homology-directed recombination templates

Knock-in experiments were performed as previously described for AML and U2OS cell lines ^35^. In brief, synthetic guide RNAs (sgRNAs) targeting the wild-type and mutant NPM1 alleles (sgNPM1wt and sgNPM1c, respectively) were purchased from Synthego (**Table S1**). DNA templates encoding fluorescent proteins muGFP and mCherry were purchased from Addgene and Twist Biosciences respectively. Templates were PCR amplified with KAPA HiFI HotStart ReadyMix (Roche) and custom oligomer primers to add flanking 100-200bp homology arms, generating locus-specific homologous recombination repair templates (**Table S1**). PCR products were purified using AMPure XP beads (Beckman Coulter) following manufacturer’s instructions.

To generate fusion proteins, stop codons were replaced with a GSG linker followed by the fluorescent-protein-coding sequence and a stop codon.

#### Cas9-sgRNA Pre-complexing and electroporation for endogenous labeling

All experiments were performed using the Neon Transfection System (ThermoFisher) as previously described^35^. Cells were prepared in Buffer R (ThermoFisher) according to manufacturer recommendations. 2.5 x 10^5^ cells were used per reaction for AML cell lines, and 6 x 10^4^ cells per reaction for U2OS cells. To obtain Cas9-sgRNA ribonucleoprotein complexes, 1 µg of Cas9 protein (PNA bio) was incubated with 1 µg of sgRNA (Synthego) for 20 minutes at room temperature. For knock-in of endogenous fluorescent labels,1 µg of homology-directed recombination template was added to each reaction. AML cell lines were electroporated at the following parameters: 1400 V, 10 ms, 3 pulses. U2OS cells were electroporated at the following parameters: 1350 V, 35 ms, 1 pulse.

For PDX2 and primary CD34+ cells, 2.5 x 10^5^ cells were prepared in Buffer T (ThermoFisher) for each reaction, and Cas9-sgRNA complexing was performed as described above. These cells were electroporated at the following parameters: 1400 V, 10 ms, 3 pulses.

### Method details

#### High-Resolution Microscopy

All imaging was performed on a Nikon Ti2E equipt with a VisiTech instant SIM microscope (VT-iSIM) with a Hamamatsu ORCA Quest qCMOS camera (C15550-20UP) camera using a CFI60 Plan Apochromat Lambda D 100X oil immersion objective lens (N.A. 1.45). Imaging conditions (i.e. laser intensity) were optimized to increase signal to noise within the dynamic range. Proteins tagged with GFP were imaged using a 488nm laser, mCherry with 561nm, and Alexa 647 with 642nm. In instances where two separate fluorophores are imaged concurrently, sequential scanning was used.

#### Live Cell Microscopy

All imaging was performed on glass-bottom 96-well plates (Cellvis P96-1.5H-N) or 8-well glass-bottom chamber slides (Ibidi #80826) coated with Poly-D-Lysine (Thermofisher, A3890401) or fibronectin (10ng/mL) for suspension and adherent cell lines respectively. Cells were maintained in 5% CO2 at 37°C during all live imaging sessions. For live cell imaging of chromatin, SPY650-DNA dye (Cytoskeleton, Inc, SC501) was added to media at a 1000X dilution according to manufacturer instructions and cells were incubated with dye for 1 hour at 37°C before imaging.

#### Immunostaining of human cell lines, PDX models, and primary patient samples

96-well glass bottom plates were incubated with poly-D-lysine (ThermoFisher, A3890401) at 37°C for 15 minutes. Following incubation, wells were aspirated and seeded with 1-2.5 x 10^5^ cells per well. The cells were incubated at 37°C for 15 minutes to allow adherence, then fixed with 4% paraformaldehyde (Sigma-Aldrich, F8775-25ML) in PBS (GenDepot, CA008-300) for 10 minutes at room temperature and washed with cold PBS three times. Fixed cells were simultaneously permeabilized and blocked with 0.5% Triton-X-100 (Sigma-Aldrich, 93443-100ML), 5% normal goat serum (ThermoFisher, 50062Z), 2% BSA (GenDepot, A0100-010), 0.1% Tween-20 (BioRad, 170-6531) in PBS at room temperature for 1 hour. Cells were then stained with rabbit or mouse primary antibodies in working solution (5% NGS, 2% BSA, 0.1% Tween-20 in PBS at 4°C overnight. Cells were washed 3 times with working solution before incubation with fluorescent secondary antibodies (Goat anti-rabbit or anti-mouse IgG, Alexa-Fluor 594/488 (ThermoFisher A-11012, A-11005,A-11029,A-11008) or Donkey Anti-Rabbit, Alexa-Fluor 647 (Abcam ab150075)) in working solution for 1 hour at room temperature. Finally, cells were washed 3 times with cold working solution, stored in PBS, and protected from light, at 4°C, until imaging.

#### Cut-and-run Sequencing

Cut-and-run sequencing was performed using Epicypher® CUTANA™ CUT&RUN protocol (V2.0) as described previously ^81^.100,000 cells were harvested per replicate per sample. Primary antibodies (anti-CRM1, anti-NUP98, anti-MLL, anti-MENIN, anti-H3K27me3, and IgG) were diluted 1:100 for overnight incubation, and DNA was purified with CUTANA™ DNA Purification Kit (Epicypher, 14-0050) according to the manufacturer’s protocol. Cut-and-run sequencing library preparation was performed using NEBNext® Ultra™ II DNA Library Prep Kit for Illumina (New England Biolabs, E7645) according to the manufacturer’s protocol. Library quality was assessed with TapeStation D5000 ScreenTape (Agilent, 5067-5588). Libraries were then sequenced using an Illumina NovaSeq 6000 sequencer, aiming for >10 million reads per condition. Paired-end enriched DNA-sequencing reads were obtained.

#### Pharmacological disruption of condensate formation and composition

To induce degradation of NPM1c, OCI-AML3 degron cells were treated with 500nM dTAG-13 (Tocris, 6605) for 24hrs. NPM1c protein degradation was confirmed via western blotting, flow-cytometry, and live-cell microscopy. To inhibit XPO1-NPM1c interactions, cells were treated with 100nM Eltanexor (Selleckchem, S8397) or 100nM Selinexor (Selleckchem, S7252) for 24hrs as previously described ^53^. To inhibit the KMT2A-MENIN interaction, cells were treated with 300nM VTP50469 (Selleckchem, S8934) or 2.5μM MI-503 (Selleckchem, S7817). 0.05% DMSO was used as a control treatment in all relevant experiments.

#### Lentiviral Plasmid Generation for Truncation Screen

Custom pLV-bsd-EF1A>(XbaI)NPM1c-mCherry-NES(BamHI) lentiviral vector, herein pLV, was purchased from VectorBuilder. To generate individual plasmids for truncation screen, pLV was linearized with XbaI (NEB, R0145S) and BamHI-HF (NEB, R3136S), purified using QIAquick Gel Extraction Kit (Qiagen, 28704, and treated with Quick-CIP (NEB, M0525S) to prevent re-ligation.

DNA oligos encoding each truncation (Twist Biosciences, **Table S1**) were cloned into linearized pLV through 2-fragment assembly using NEBuilder^®^ HiFi DNA Assembly Master Mix (NEB, E2621S) according to manufacturer recommendations. Assembled plasmids were transformed into TOP10 competent cells and selected on LB plates with Ampicillin (50 mg/ml) (Sigma-Aldrich, A5354-10ML) overnight before single colony isolation, plasmid extraction and validation with whole-plasmid long-read sequencing (Plasmidsaurus).

#### Lentivirus Generation for Truncation Screen

6-well dishes of 80-90% confluent Lenti-X 293T cells were transfected using Lipofectamine 3000 (Invitrogen), 1.25µg of lentiviral plasmid, 625ng of pMD2.G., and 625ng of pCMV delta R8.2 (packaging vectors). Media containing lentiviral particles was collected at 48h and 72h after transfection. To concentrate viral particles, viral media was incubated with 4X polyethylene glycol (PEG) (32% PEG6000, 0.4M NaCl, and 0.04M 4-(2-hydroxyethyl)-1-piperazineethanesulfonic acid (HEPES)) while shaking at 4°C overnight. The solution was then centrifuged at 1500xg for 45 minutes and resuspended in RPMI supplemented with 10% FBS in 1% of the original volume. Concentrated viral aliquots were stored at -80°C.

#### NPM1c Truncation Screen

2.5 x 10^5^ OCI-AML3 degron cells were seeded in 24-well plates with RPMI supplemented with polybrene (7μg/ml) and 2.5μl of concentrated viral particles. Cells were spin infected at 1100rpm for 2 hrs at room temperature, and returned to the incubator overnight. 24hrs after transduction, cells were washed once with PBS and replaced with fresh media. 48hrs after transduction, cells were treated with 500nM dTAG-13 or DMSO. Cells were split 1:6 every 3 days, and fresh media with 500nM dTAG-13 or DMSO was added. Flow cytometry was performed every 2-3 days to assess mCherry expression within the population. Live cell imaging was performed at day 7 as described in methods (this paper).

Flow cytometry for myeloid differentiation markers was also performed on day 14 as described previously ^35^. Briefly, cells were pelleted at 300g for 5 minutes and resuspended in 50μL flow buffer (25mM HEPES, 1mM EDTA pH 8, 3% FBS in PBS) with anti-CD11b-Pacific Blue antibodies (1:100) (Biolegend, 101223). Cells were incubated at room temperature in the dark for 15 minutes and 300μL flow buffer was added. Cells were gently vortexed before pelleting. Supernatant was removed, and cells resuspended in 150μL flow buffer. All samples were covered and stored on ice until analyzed. Single color controls (cell lines and UltraComp eBeads (Thermofisher, 01-3333-42) were utilized for all flow cytometry experiments for minimal compensation. All samples were run on a BD LSRII benchtop cytometer.

#### Transfection of U2OS cells

U2OS cells were seeded at 2x10^4^/mL in a 96-well glass-bottom dish coated with fibronectin (10ng/mL), and placed in an incubator overnight to allow adherence. On the following day, cells were transfected with Lipofectamine 3000 (Thermofisher, L3000015) according to manufacturer recommendations, and 100-300ng plasmid DNA. Media was replaced the next day, and cells were imaged 48hrs after transfection.

#### DNA FISH

DNA FISH protocol was adapted from Liu *et al*^82^. Briefly, OCI-AML3 NPM1c-muGFP cells were fixed with 4% PFA, permeabilized with Triton X-100 and stained with anti-NUP98 antibodies. After staining with the secondary antibodies, cells were cross-linked with 2% PFA and incubated with the anti-*HOXA9* FISH probe (HOXA9-20-O-R, Empire Genomics).

#### *Ex vivo* VTP50469 treatment of murine HSPCs

Bone marrow cells from mice with >80% GFP+ cells in the peripheral blood and bone marrow were isolated and plated 100,000 cells per well in U-bottom 96-well plates in PVA-based culture media. Cells were treated with 300 nM VTP50469 or vehicle for 72h (qRT-PCR analysis) or 6 days (flow cytometry analysis).

#### Flow cytometry analysis

Murine cells were stained with CD117, Mac1, CD4, CD8, B220 antibodies, and Sytox Blue (viability dye) and analyzed with BD Fortessa.

#### qPCR analysis

1000 ng of RNA was isolated using RNAeasy Mini Kit (Qiagen), treated with DNAse, and used for reverse transcription reaction (RevertAid RT Reverse Transcription Kit, ThermoFisher Scientific). qRT-PCR reaction was performed using the following primers: *m_meis1*: TATGTGACAATTTCTGCCACCG, AGTGGATGCCGTGTCATCAT; *m_hoxa9*: TGTCCCACGCTTGACACTC, AGCGAGCATGTAGCCAGTTG; *m_hoxa10*: CGAGTCCTAGACTCCACGC, CAGTTGGCTGCATTTTCGCC.

#### Lentiviral and retroviral transduction of murine HSPCs

C-kit+ cells from murine bone marrow were purified using anti-CD117 microbeads (#130-091-224, Miltenyi Biotec) and plated 100000 cells per well 96-well plates in PVA-based culture media. Media composition was adapted from Khoo *et al*.^83^ to ensure the best recovery of cells after viral transduction. Cells were transduced with PEG-concentrated retro- (NrasG12D:GFP) and lentiviral (FL, dNES, mCherry) particles in the presence of 7 ug/ml of polybrene. 48 hours after transduction the efficacy was quantified with flow cytometry, and cells were used for a competitive bone marrow transplantation (1:10 ratio) into wild-type C57BL/6 recipient mice. The percentage of GFP or mCherry cells in the peripheral blood was monitored bi-weekly.

#### Immunoblotting

Cells were lysed with RIPA lysis buffer supplemented with the protease inhibitor. 10-15 ug of the total protein were separated by SDS-PAGE, transferred to the nitrocellulose membranes (0.45 µm), and probed using anti-GAPDH (Rhodamine-conjugated, #12004168, Bio-Rad), anti-NPM1wt (NB600-1030, Novus Biologicals), anti-NPM1c (NB110-61646, Novus Biologicals) and anti-GFP (NB600-308, Novus Biologicals) antibodies. We used fluorescent secondary antibodies (anti-rabbit StarBright 700 and anti-mouse StarBright 520, Bio-Rad) to detect the proteins using the Bio-Rad Chemidoc imaging system.

#### Selection and determination of fusion protein sequences

Gene fusions were identified by computational (TopHat-Fusion) analysis of transcriptome sequencing data from the Cancer Genome Atlas (TCGA) (RRID:SCR_003193). To obtain fusion proteins, the list was filtered for “in-frame” gene fusions (*i.e.*, chimeric proteins made from joining two genes that usually encode separate proteins). Three fusion oncoproteins (NUP98::NSD1, KMT2A::AFF1, and SET::NUP214) were selected for this study. Their breakpoints or junctions were prioritized based on the highest frequencies of occurrence found in TCGA: NUP98 (chr11:3744509) and NSD1 (chr5:177235821); KMT2A (chr11:118482495) and AFF1 (chr4:87089984); SET (chr9:128694042) and NUP214 (chr9:131159383).

### Quantification and Statistical Analysis

#### Quantitative Microscopy

All imaging data was analyzed in Micro-manager using a custom-built imaging plugin adapted from AcqEngJ and NDViewer. Roughly every other day, power meter measurements were done at the focal plane to verify microscope performance and allow for day-to-day comparison of measurements. Roughly every month, dye standards were used to calibrate the DL offset and DL to photon conversion by fitting the histogram of photon counts at lower light, produce a conversion table to reference settings, and background and flat field corrections functions (functions because it was dependent on laser intensity). Linearity of the camera was verified manually as needed. For images, background field, flat field corrections, and conversion table were applied were used. Critically during quantification of intensity within an ROIs, average values within the ROI on the image background field subtracted and the flat field were calculated separately and then divided.

#### ROI determination in live or fixed cells

For all quantification, representative areas corresponding to dilute phases of individual cellular compartments (e.g. cytoplasm and nucleoplasm) were manually selected per cell. Representative areas of dense phase structures (e.g. nucleoli and nuclear condensates) were find max over an ROI (e.g. nucleus or cell) defined by a box of size 3 by 3 pixels (when appropriate larger sizes were used).

#### Quantification of brightest puncta enrichment and radial distribution function in live or fixed cells

Regions of interest (ROIs) were first selected by manually drawn polygon outlines within nuclei of cells for analysis. Individuals drawing nuclear masks were blinded to fluorescent channel under investigation for enrichment in the nucleus (e.g. only NPM1c was observed when quantifying XPO1 enrichment in nuclear puncta). Radial Distribution Function calculation was performed as previously described^50^ but with flat field measurements included in the denominator of the RDF and the routine being adapted into the custom built gui within micromanager.

#### Fitting microscopy data

The distribution for condensates/no condensates (Figure 4 & 6) as a function of concentration was determined by finding the maximum likelihood values for the sigmoidal dependence. Partitioning values were extrapolated by fitting the dilute and dense concentrations with a hyperbola and using the instantaneous slope as the partitioning. An iterative weighting routine was used to remove outliers using a protocol adapted from ^44^.

#### Cut-and-run Sequencing Analysis

CUT-and-run Sequencing analysis was performed as previously described^81^. In brief, paired-end sequencing data were mapped to the human genome (hg19) using Bowtie2. Reads from biological replicates were then merged for peak calling using MACS3. Quantification of peaks was performed using Subread package, where peak regions were identified as 3kb regions centered around transcriptional start sites.

#### RNA Sequencing

Raw paired-end sequencing data was preprocessed using Fastp to exclude: (i) read lengths < 30, (ii) low quality reads (Q<25), (iii) adapter sequences. Filtered reads were mapped to the human reference (hg38) using STAR. Gene-level quantification was performed through featureCounts. Low expressed genes (<10 counts) were removed before normalizing with DESeq2.

## References

1. Lyon, A.S., Peeples, W.B., and Rosen, M.K. (2021). A framework for understanding the functions of biomolecular condensates across scales. Nat. Rev. Mol. Cell Biol. 22, 215– 235.

2. Grisendi, S., Mecucci, C., Falini, B., and Pandolfi, P.P. (7/2006). Nucleophosmin and cancer. Nat. Rev. Cancer 6, 493–505.

3. Mitrea, D.M., Cika, J.A., Guy, C.S., Ban, D., Banerjee, P.R., Stanley, C.B., Nourse, A., Deniz, A.A., and Kriwacki, R.W. (2016). Nucleophosmin integrates within the nucleolus via multi-modal interactions with proteins displaying R-rich linear motifs and rRNA. Elife 5. 10.7554/eLife.13571.

4. Lindström, M.S. (2011). NPM1/B23: A Multifunctional Chaperone in Ribosome Biogenesis and Chromatin Remodeling. Biochem. Res. Int. 2011, 195209.

5. Yao, Z., Duan, S., Hou, D., Wang, W., Wang, G., Liu, Y., Wen, L., and Wu, M. (2010). B23 acts as a nucleolar stress sensor and promotes cell survival through its dynamic interaction with hnRNPU and hnRNPA1. Oncogene 29, 1821–1834.

6. Yang, K., Wang, M., Zhao, Y., Sun, X., Yang, Y., Li, X., Zhou, A., Chu, H., Zhou, H., Xu, J., et al. (2016). A redox mechanism underlying nucleolar stress sensing by nucleophosmin. Nat. Commun. 7, 13599.

7. Wang, X., and Li, S. (2014). Protein mislocalization: mechanisms, functions and clinical applications in cancer. Biochim. Biophys. Acta 1846, 13–25.

8. Koken, M.H., Puvion-Dutilleul, F., Guillemin, M.C., Viron, A., Linares-Cruz, G., Stuurman, N., de Jong, L., Szostecki, C., Calvo, F., and Chomienne, C. (1994). The t(15;17) translocation alters a nuclear body in a retinoic acid-reversible fashion. EMBO J. 13, 1073– 1083.

9. de Thé, Hugues, Le Bras, M., and Lallemand-Breitenbach, V. (2012). Acute promyelocytic leukemia, arsenic, and PML bodies. J. Cell Biol. 198, 11–21.

10. Banani, S.F., Lee, H.O., Hyman, A.A., and Rosen, M.K. (2017). Biomolecular condensates: organizers of cellular biochemistry. Nat. Rev. Mol. Cell Biol. 18, 285–298.

11. Hnisz, D., Shrinivas, K., Young, R.A., Chakraborty, A.K., and Sharp, P.A. (2017). A Phase Separation Model for Transcriptional Control. Cell 169, 13–23.

12. Rosencrance, C.D., Ammouri, H.N., Yu, Q., Ge, T., Rendleman, E.J., Marshall, S.A., and Eagen, K.P. (2020). Chromatin Hyperacetylation Impacts Chromosome Folding by Forming a Nuclear Subcompartment. Mol. Cell 78, 112–126.e12.

13. Kosno, M., Currie, S.L., Kumar, A., Xing, C., and Rosen, M.K. (2023). Molecular features driving condensate formation and gene expression by the BRD4-NUT fusion oncoprotein are overlapping but distinct. Sci. Rep. 13, 11907.

14. Song, L., Yao, X., Li, H., Peng, B., Boka, A.P., Liu, Y., Chen, G., Liu, Z., Mathias, K.M., Xia, L., et al. (2022). Hotspot mutations in the structured ENL YEATS domain link aberrant transcriptional condensates and cancer. Mol. Cell 0, 4080–4098.e12.

15. Chandra, B., Michmerhuizen, N.L., Shirnekhi, H.K., Tripathi, S., Pioso, B.J., Baggett, D.W., Mitrea, D.M., Iacobucci, I., White, M.R., Chen, J., et al. (2022). Phase Separation Mediates NUP98 Fusion Oncoprotein Leukemic Transformation. Cancer Discov. 12, 1152–1169.

16. Terlecki-Zaniewicz, S., Humer, T., Eder, T., Schmoellerl, J., Heyes, E., Manhart, G., Kuchynka, N., Parapatics, K., Liberante, F.G., Müller, A.C., et al. (2021). Biomolecular condensation of NUP98 fusion proteins drives leukemogenic gene expression. Nat. Struct. Mol. Biol. 28, 190–201.

17. Banani, S.F., Afeyan, L.K., Hawken, S.W., Henninger, J.E., Dall’Agnese, A., Clark, V.E., Platt, J.M., Oksuz, O., Hannett, N.M., Sagi, I., et al. (2022). Genetic variation associated with condensate dysregulation in disease. Dev. Cell 57, 1776–1788.e8.

18. Tripathi, S., Shirnekhi, H.K., Gorman, S.D., Chandra, B., Baggett, D.W., Park, C.-G., Somjee, R., Lang, B., Hosseini, S.M.H., Pioso, B.J., et al. (2023). Defining the condensate landscape of fusion oncoproteins. Nat. Commun. 14, 6008.

19. Yoo, H., Triandafillou, C., and Drummond, D.A. (2019). Cellular sensing by phase separation: Using the process, not just the products. J. Biol. Chem. 294, 7151–7159.

20. Mittag, T., and Pappu, R.V. (2022). A conceptual framework for understanding phase separation and addressing open questions and challenges. Mol. Cell 82, 2201–2214.

21. Pappu, R.V., Cohen, S.R., Dar, F., Farag, M., and Kar, M. (2023). Phase Transitions of Associative Biomacromolecules. Chem. Rev. 123, 8945–8987.

22. Riback, J.A., Zhu, L., Ferrolino, M.C., Tolbert, M., Mitrea, D.M., Sanders, D.W., Wei, M.-T., Kriwacki, R.W., and Brangwynne, C.P. (2020). Composition-dependent thermodynamics of intracellular phase separation. Nature 581, 209–214.

23. Li, P., Banjade, S., Cheng, H.-C., Kim, S., Chen, B., Guo, L., Llaguno, M., Hollingsworth, J.V., King, D.S., Banani, S.F., et al. (2012). Phase transitions in the assembly of multivalent signalling proteins. Nature 483, 336–340.

24. Sanders, D.W., Kedersha, N., Lee, D.S.W., Strom, A.R., Drake, V., Riback, J.A., Bracha, D., Eeftens, J.M., Iwanicki, A., Wang, A., et al. (2020). Competing Protein-RNA Interaction Networks Control Multiphase Intracellular Organization. Cell 181, 306–324.e28.

25. Mitrea, D.M., Grace, C.R., Buljan, M., Yun, M.-K., Pytel, N.J., Satumba, J., Nourse, A., Park, C.-G., Madan Babu, M., White, S.W., et al. (2014). Structural polymorphism in the N-terminal oligomerization domain of NPM1. Proc. Natl. Acad. Sci. U. S. A. 111, 4466–4471.

26. Mitrea, D.M., Cika, J.A., Stanley, C.B., Nourse, A., Onuchic, P.L., Banerjee, P.R., Phillips, A.H., Park, C.G., Deniz, A.A., and Kriwacki, R.W. (2018). Self-interaction of NPM1 modulates multiple mechanisms of liquid-liquid phase separation. Nat. Commun. 9. 10.1038/s41467-018-03255-3.

27. Falini, B., Mecucci, C., Tiacci, E., Alcalay, M., Rosati, R., Pasqualucci, L., La Starza, R., Diverio, D., Colombo, E., Santucci, A., et al. (2005). Cytoplasmic nucleophosmin in acute myelogenous leukemia with a normal karyotype. N. Engl. J. Med. 352, 254–266.

28. Schlenk, R.F., Döhner, K., Krauter, J., Fröhling, S., Corbacioglu, A., Bullinger, L., Habdank, M., Späth, D., Morgan, M., Benner, A., et al. (2008). Mutations and treatment outcome in cytogenetically normal acute myeloid leukemia. N. Engl. J. Med. 358, 1909–1918.

29. Bolli, N., Nicoletti, I., De Marco, M.F., Bigerna, B., Pucciarini, A., Mannucci, R., Martelli, M.P., Liso, A., Mecucci, C., Fabbiano, F., et al. (2007). Born to be exported: COOH-terminal nuclear export signals of different strength ensure cytoplasmic accumulation of nucleophosmin leukemic mutants. Cancer Res. 67, 6230–6237.

30. Grummitt, C.G., Townsley, F.M., Johnson, C.M., Warren, A.J., and Bycroft, M. (2008). Structural consequences of nucleophosmin mutations in acute myeloid leukemia. J. Biol. Chem. 283, 23326–23332.

31. Falini, B., Bolli, N., Shan, J., Martelli, M.P., Liso, A., Pucciarini, A., Bigerna, B., Pasqualucci, L., Mannucci, R., Rosati, R., et al. (2006). Both carboxy-terminus NES motif and mutated tryptophan(s) are crucial for aberrant nuclear export of nucleophosmin leukemic mutants in NPMc+ AML. Blood 107, 4514–4523.

32. Azmi, A.S., Uddin, M.H., and Mohammad, R.M. (2020). The nuclear export protein XPO1 — from biology to targeted therapy. Nat. Rev. Clin. Oncol. 18, 152–169.

33. Alcalay, M., Tiacci, E., Bergomas, R., Bigerna, B., Venturini, E., Minardi, S.P., Meani, N., Diverio, D., Bernard, L., Tizzoni, L., et al. (2005). Acute myeloid leukemia bearing cytoplasmic nucleophosmin (NPMc+ AML) shows a distinct gene expression profile characterized by up-regulation of genes involved in stem-cell maintenance. Blood 106, 899–902.

34. Spencer, D.H., Young, M.A., Lamprecht, T.L., Helton, N.M., Fulton, R., O’Laughlin, M., Fronick, C., Magrini, V., Demeter, R.T., Miller, C.A., et al. (2015). Epigenomic analysis of the HOX gene loci reveals mechanisms that may control canonical expression patterns in AML and normal hematopoietic cells. Leukemia 29, 1279–1289.

35. Brunetti, L., Gundry, M.C., Sorcini, D., Guzman, A.G., Huang, Y.H., Ramabadran, R., Gionfriddo, I., Mezzasoma, F., Milano, F., Nabet, B., et al. (2018). Mutant NPM1 Maintains the Leukemic State through HOX Expression. Cancer Cell 34, 499–512.e9.

36. Uckelmann, H.J., Haarer, E.L., Takeda, R., Wong, E.M., Hatton, C., Marinaccio, C., Perner, F., Rajput, M., Antonissen, N.J.C., Wen, Y., et al. (2023). Mutant NPM1 Directly Regulates Oncogenic Transcription in Acute Myeloid Leukemia. Cancer Discov. 13, 746–765.

37. Wang, X.Q.D., Fan, D., Han, Q., Liu, Y., Miao, H., Wang, X., Li, Q., Chen, D., Gore, H., Himadewi, P., et al. (2023). Mutant NPM1 Hijacks Transcriptional Hubs to Maintain Pathogenic Gene Programs in Acute Myeloid Leukemia. Cancer Discov. 13, 724–745.

38. Falini, B. (2023). NPM1-mutated acute myeloid leukemia: New pathogenetic and therapeutic insights and open questions. Am. J. Hematol. 98, 1452–1464.

39. Falini, B., Brunetti, L., Sportoletti, P., and Martelli, M.P. (2020). NPM1-mutated acute myeloid leukemia: from bench to bedside. Blood 136, 1707–1721.

40. Heikamp, E.B., Henrich, J.A., Perner, F., Wong, E.M., Hatton, C., Wen, Y., Barwe, S.P., Gopalakrishnapillai, A., Xu, H., Uckelmann, H.J., et al. (2022). The menin-MLL1 interaction is a molecular dependency in NUP98-rearranged AML. Blood 139, 894–906.

41. Issa, G.C., Aldoss, I., DiPersio, J., Cuglievan, B., Stone, R., Arellano, M., Thirman, M.J., Patel, M.R., Dickens, D.S., Shenoy, S., et al. (2023). The menin inhibitor revumenib in KMT2A-rearranged or NPM1-mutant leukaemia. Nature 615, 920–924.

42. Uckelmann, H.J., Kim, S.M., Wong, E.M., Hatton, C., Giovinazzo, H., Gadrey, J.Y., Krivtsov, A.V., Rücker, F.G., Döhner, K., Mcgeehan, G.M., et al. (2020). Therapeutic targeting of preleukemia cells in a mouse model of NPM1 mutant acute myeloid leukemia. 590, 586–590.

43. Mendes, A., Jühlen, R., Martinelli, V., and Fahrenkrog, B. (2020). Targeted CRM1-inhibition perturbs leukemogenic NUP214 fusion proteins and exerts anti-cancer effects in leukemia cell lines with NUP214 rearrangements. Oncotarget 11, 3371–3386.

44. Dollinger, C., Potolitsyna, E., Martin, A.G., Anand, A., Datar, G.K., Schmit, J.D., and Riback, J.A. (2025). Nanometer condensate organization in live cells derived from partitioning measurements. bioRxivorg. 10.1101/2025.02.26.640428.

45. Hernandez-Verdun, D. (05/2011). Assembly and disassembly of the nucleolus during the cell cycle. Nucleus 2, 189–194.

46. Lafontaine, D.L.J., Riback, J.A., Bascetin, R., and Brangwynne, C.P. (03/2021). The nucleolus as a multiphase liquid condensate. Nat. Rev. Mol. Cell Biol. 22, 165–182.

47. Rai, A.K., Chen, J.-X., Selbach, M., and Pelkmans, L. (2018). Kinase-controlled phase transition of membraneless organelles in mitosis. Nature 559, 211–216.

48. Güttinger, S., Laurell, E., and Kutay, U. (3/2009). Orchestrating nuclear envelope disassembly and reassembly during mitosis. Nat. Rev. Mol. Cell Biol. 10, 178–191.

49. Wu, Z., Jiang, Q., Clarke, P.R., and Zhang, C. (2013). Phosphorylation of Crm1 by CDK1-cyclin-B promotes Ran-dependent mitotic spindle assembly. J. Cell Sci. 126, 3417–3428.

50. Riback, J.A., Eeftens, J.M., Lee, D.S.W., Quinodoz, S.A., Donlic, A., Orlovsky, N., Wiesner, L., Beckers, L., Becker, L.A., Strom, A.R., et al. (2023). Viscoelasticity and advective flow of RNA underlies nucleolar form and function. Mol. Cell 83, 3095–3107.e9.

51. Matkar, S., Thiel, A., and Hua, X. (2013). Menin: a scaffold protein that controls gene expression and cell signaling. Trends Biochem. Sci. 38, 394–402.

52. Hing, Z.A., Fung, H.Y.J., Ranganathan, P., Mitchell, S., El-Gamal, D., Woyach, J.A., Williams, K., Goettl, V.M., Smith, J., Yu, X., et al. (2016). Next-generation XPO1 inhibitor shows improved efficacy and in vivo tolerability in hematological malignancies. Leukemia 30, 2364–2372.

53. Pianigiani, G., Gagliardi, A., Mezzasoma, F., Rocchio, F., Tini, V., Bigerna, B., Sportoletti, P., Caruso, S., Marra, A., Peruzzi, S., et al. (2022). Prolonged XPO1 inhibition is essential for optimal antileukemic activity in NPM1-mutated AML. Blood Adv 6, 5938–5949.

54. Garzon, R., Savona, M., Baz, R., Andreeff, M., Gabrail, N., Gutierrez, M., Savoie, L., Mau-Sorensen, P.M., Wagner-Johnston, N., Yee, K., et al. (2017). A phase 1 clinical trial of single-agent selinexor in acute myeloid leukemia. Blood 129, 3165–3174.

55. Oka, M., Mura, S., Otani, M., Miyamoto, Y., Nogami, J., Maehara, K., Harada, A., Tachibana, T., Yoneda, Y., and Ohkawa, Y. (2019). Chromatin-bound CRM1 recruits SET-Nup214 and NPM1c onto HOX clusters causing aberrant HOX expression in leukemia cells. Elife 8. 10.7554/eLife.46667.

56. Krivtsov, A.V., Evans, K., Gadrey, J.Y., Eschle, B.K., Hatton, C., Uckelmann, H.J., Ross, K.N., Perner, F., Olsen, S.N., Pritchard, T., et al. (2019). A Menin-MLL Inhibitor Induces Specific Chromatin Changes and Eradicates Disease in Models of MLL-Rearranged Leukemia. Cancer Cell 36, 660–673.e11.

57. Loberg, M.A., Bell, R.K., Goodwin, L.O., Eudy, E., Miles, L.A., SanMiguel, J.M., Young, K., Bergstrom, D.E., Levine, R.L., Schneider, R.K., et al. (2019). Sequentially inducible mouse models reveal that Npm1 mutation causes malignant transformation of Dnmt3a-mutant clonal hematopoiesis. Leukemia 33, 1635–1649.

58. Dovey, O.M., Cooper, J.L., Mupo, A., Grove, C.S., Lynn, C., Conte, N., Andrews, R.M., Pacharne, S., Tzelepis, K., Vijayabaskar, M.S., et al. (2017). Molecular synergy underlies the co-occurrence patterns and phenotype of NPM1-mutant acute myeloid leukemia. Blood 130, 1911–1922.

59. Hingorani, K., Szebeni, A., and Olson, M.O. (2000). Mapping the functional domains of nucleolar protein B23. J. Biol. Chem. 275, 24451–24457.

60. Michmerhuizen, N.L., Klco, J.M., and Mullighan, C.G. (2020). Mechanistic insights and potential therapeutic approaches for NUP98-rearranged hematologic malignancies. Blood 136, 2275–2289.

61. Gough, S.M., Slape, C.I., and Aplan, P.D. (2011). NUP98 gene fusions and hematopoietic malignancies: common themes and new biologic insights. Blood 118, 6247–6257.

62. Meyer, C., Larghero, P., Almeida Lopes, B., Burmeister, T., Gröger, D., Sutton, R., Venn, N.C., Cazzaniga, G., Corral Abascal, L., Tsaur, G., et al. (2023). The KMT2A recombinome of acute leukemias in 2023. Leukemia 37, 988–1005.

63. Oka, M., Otani, M., Miyamoto, Y., Oshima, R., Adachi, J., Tomonaga, T., Asally, M., Nagaoka, Y., Tanaka, K., Toyoda, A., et al. (08/2023). Phase-separated nuclear bodies of nucleoporin fusions promote condensation of MLL1/CRM1 and rearrangement of 3D genome structure. Cell Rep. 42, 112884.

64. Feric, M., Vaidya, N., Harmon, T.S., Mitrea, D.M., Zhu, L., Richardson, T.M., Kriwacki, R.W., Pappu, R.V., and Brangwynne, C.P. (2016). Coexisting Liquid Phases Underlie Nucleolar Subcompartments. Cell 165, 1686–1697.

65. Trinkle-Mulcahy, L., and Sleeman, J.E. (2017). The Cajal body and the nucleolus: “In a relationship” or “It’s complicated”? RNA Biol. 14, 739–751.

66. Gu, X., Ebrahem, Q., Mahfouz, R.Z., Hasipek, M., Enane, F., Radivoyevitch, T., Rapin, N., Przychodzen, B., Hu, Z., Balusu, R., et al. (2018). Leukemogenic nucleophosmin mutation disrupts the transcription factor hub that regulates granulomonocytic fates. J. Clin. Invest. 128, 4260–4279.

67. Wang, A.J., Han, Y., Jia, N., Chen, P., and Minden, M.D. (2020). NPM1c impedes CTCF functions through cytoplasmic mislocalization in acute myeloid leukemia. Leukemia 34, 1278–1290.

68. Wu, H.-C., Rerolle, D., Berthier, C., Hleihel, R., Sakamoto, T., Quentin, S., Benhenda, S., Morganti, C., Wu, C., Conte, L., et al. (2021). Actinomycin D targets NPM1c-primed mitochondria to restore PML-driven senescence in AML therapy. Cancer Discov., candisc.0177.2021.

69. Gourvest, M., De Clara, E., Wu, H.-C., Touriol, C., Meggetto, F., De Thé, H., Pyronnet, S., Brousset, P., and Bousquet, M. (2021). A novel leukemic route of mutant NPM1 through nuclear import of the overexpressed long noncoding RNA LONA. Leukemia 35, 2784–2798.

70. Sumner, M.C., and Brickner, J. (2022). The Nuclear Pore Complex as a Transcription Regulator. Cold Spring Harb. Perspect. Biol. 14. 10.1101/cshperspect.a039438.

71. Ranganathan, P., Yu, X., Na, C., Santhanam, R., Shacham, S., Kauffman, M., Walker, A., Klisovic, R., Blum, W., Caligiuri, M., et al. (2012). Preclinical activity of a novel CRM1 inhibitor in acute myeloid leukemia. Blood 120, 1765–1773.

72. Grisendi, S., Bernardi, R., Rossi, M., Cheng, K., Khandker, L., Manova, K., and Pandolfi, P.P. (2005). Role of nucleophosmin in embryonic development and tumorigenesis. Nature 437, 147–153.

73. Shlush, L.I., Zandi, S., Mitchell, A., Chen, W.C., Brandwein, J.M., Gupta, V., Kennedy, J.A., Schimmer, A.D., Schuh, A.C., Yee, K.W., et al. (2014). Identification of pre-leukaemic haematopoietic stem cells in acute leukaemia. Nature 506, 328–333.

74. Cancer Genome Atlas Research Network, Ley, T.J., Miller, C., Ding, L., Raphael, B.J., Mungall, A.J., Robertson, A.G., Hoadley, K., Triche, T.J., Jr, Laird, P.W., et al. (2013). Genomic and epigenomic landscapes of adult de novo acute myeloid leukemia. N. Engl. J. Med. 368, 2059–2074.

75. Yamazaki, H., Takagi, M., Kosako, H., Hirano, T., and Yoshimura, S.H. (2022). Cell cycle-specific phase separation regulated by protein charge blockiness. Nat. Cell Biol. 24, 625– 632.

76. Zhou, M.-H., and Yang, Q.-M. (2014). NUP214 fusion genes in acute leukemia (Review). Oncol. Lett. 8, 959–962.

77. Winters, A.C., and Bernt, K.M. (2017). MLL-Rearranged Leukemias-An Update on Science and Clinical Approaches. Front Pediatr 5, 4.

78. Alexander, T.B., and Mullighan, C.G. (2021). Molecular Biology of Childhood Leukemia. Annual Review of Cancer Biology 5, 95–117.

79. Balasubramanian, S.K., Azmi, A.S., and Maciejewski, J. (2022). Selective inhibition of nuclear export: a promising approach in the shifting treatment paradigms for hematological neoplasms. Leukemia 36, 601–612.

80. Perner, F., Stein, E.M., Wenge, D.V., Singh, S., Kim, J., Apazidis, A., Rahnamoun, H., Anand, D., Marinaccio, C., Hatton, C., et al. (2023). MEN1 mutations mediate clinical resistance to menin inhibition. Nature 615, 913–919.

81. Chen, C.-W., Zhang, L., Dutta, R., Niroula, A., Miller, P.G., Gibson, C.J., Bick, A.G., Reyes, J.M., Lee, Y.-T., Tovy, A., et al. (11/2023). SRCAP mutations drive clonal hematopoiesis through epigenetic and DNA repair dysregulation. Cell Stem Cell 30, 1503–1519.e8.

82. Liu, Y., Li, Q., Song, L., Gong, C., Tang, S., Budinich, K.A., Vanderbeck, A., Mathias, K.M., Wertheim, G.B., Nguyen, S.C., et al. (2024). Condensate-promoting ENL mutation drives tumorigenesis in vivo through dynamic regulation of histone modifications and gene expression. Cancer Discov. 14, 1522–1546.

83. Khoo, H.M., Meaker, G.A., and Wilkinson, A.C. (2023). Ex vivo expansion and genetic manipulation of mouse hematopoietic stem cells in polyvinyl alcohol-based cultures. J. Vis. Exp. 10.3791/64791.

